# Aniline Dioxygenase in *Rhodococcus ruber* R1: Insights into Skatole Degradation

**DOI:** 10.1101/2024.12.29.630347

**Authors:** S.J. Galaz, B. Saavedra, A. Zúñiga, F. González-Toro, R. Donoso

## Abstract

Skatole is an aromatic heterocyclic compound with a strong offensive odor, produced by microorganisms during the anaerobic breakdown of tryptophan. Skatole accumulation is linked to environmental and health issues. Despite its persistence and harmful effects, skatole’s biodegradation by microorganisms is poorly understood. We have recently isolated a gram-positive bacterium, *Rhodococcus ruber* R1, which uses skatole as its sole carbon and energy source. Here we report an operon consisting of 14 genes encoding aromatic oxygenase systems involved in skatole degradation in *Rhodococcus ruber* R1. Cells growing on skatole accumulate aniline transiently, indicating its role as an intermediate in the degradation pathway. We characterize six genes in this cluster that encode for an aniline dioxygenase, which converts aniline to catechol and is only activated in the presence of skatole. This gene cluster was successfully introduced into a heterologous strain enabling the full degradation of aniline and its derivatives. Phylogenetic analysis of aniline dioxygenase present in R1 strain reveals a widespread distribution of this system among bacteria, in contrast to the full skatole cluster, which is restricted to a few genera. These findings advance our understanding of the skatole degradation pathway and highlight R1’s potential for bioremediation of skatole, aniline, and related contaminants.

## INTRODUCTION

Skatole (3-methylindole) is an aromatic heterocyclic compound renowned for its unpleasant odor, which plays a significant role in the distinctive smell of animal feces^1,2^. It is produced by microorganisms during the anaerobic fermentation of tryptophan (Trp), primarily within the intestines of mammals^2,3^. Skatole synthesis begins with the transformation of Trp through several alternative pathways into indole-3-acetic acid (IAA)^4–6^. Then, the IAA is decarboxylated by the enzyme indoleacetate decarboxylase, which leads to the formation of skatole^3^. Skatole occurrence is associated with the animal industry such as pig manure, livestock farms, and processing plants^7–10^. Skatole concentrations in wastewater have been detected as high as 700 μg/L, with skatole concentrations in air ranging from 0.024 to 1.75 ng/L, while the odor threshold level has been determined to be 0.327 ng/L by an expert panel^10^. Such high concentrations can result in significant environmental contamination, posing potential health risks (Fig. 1A). Skatole pollution is linked to acute bovine pulmonary edema and emphysema and its overproduction in the intestines is related to human putrefactive dysbiosis^11,12^. Different approaches have been investigated to eliminate skatole, such as chemical scrubbing, adsorption, or biofiltration^10,13,14^, however, none of them has been applied efficiently and without side effects on the environment.

**Figure 1.**
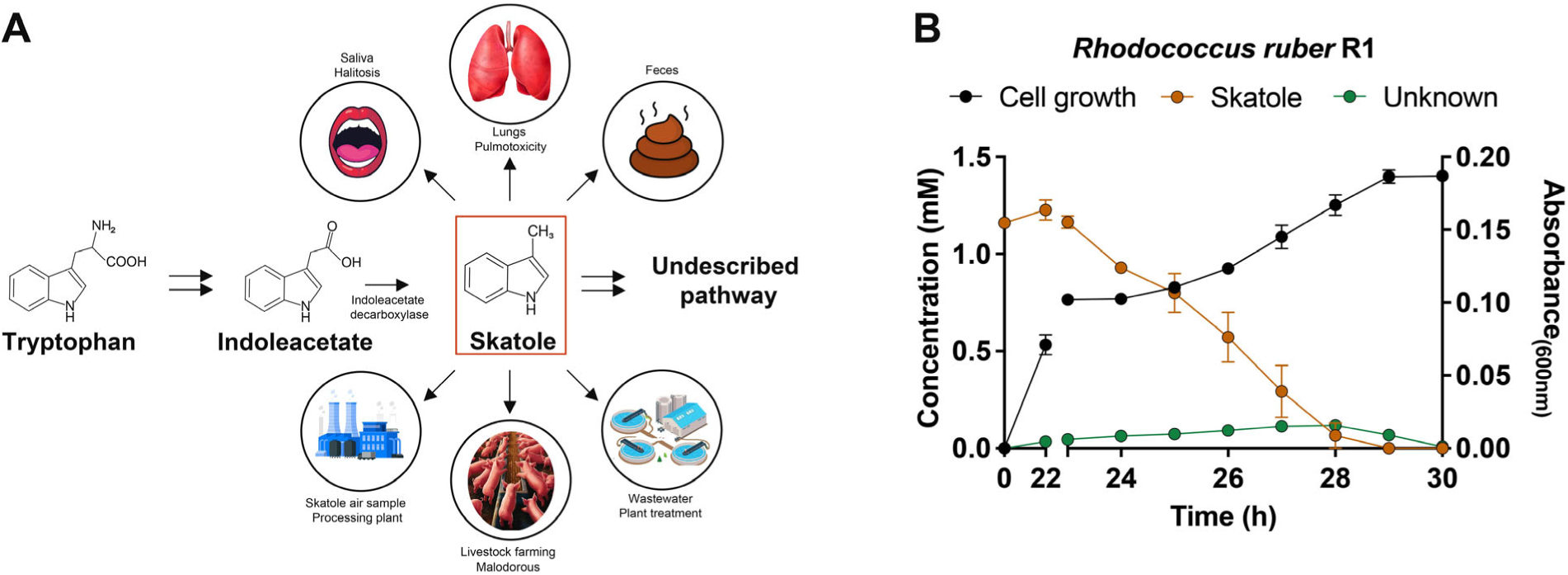
Skatole synthesis and its degradation by *Rhodococcus rube*r R1. **(A)** The tryptophan is converted into skatole via indoleacetate by the enzyme indoleacetate decarboxylase, in the intestine of mammals. Skatole is associated with halitosis, pulmonotoxicity, and putrefactive dysbiosis, and is considered an organic pollutant in processing plants, livestock farms, and wastewater. The biodegradation pathway and mechanism of skatole are not yet fully described. **(B)** Growth and degradation of skatole by strain *Rhodococcus ruber* R1. Growth curve of R1 strain on a minimal medium with 1 mM skatole as a sole carbon source (black line, absorbance at 600 nm, right axis). Skatole degradation from supernatant (orange line, skatole concentration in mM, left axis). High-performance liquid chromatography (HPLC) was performed to measure skatole degradation. An unknown intermediate is accumulated during growth (green line, concentration in mM, left axis). Experiments were performed in two biological replicates. Error bars: ±SD.

Microorganisms can overcome the limitations of skatole degradation, as they can convert or degrade it for use as a carbon source. Proteobacteria have been identified as skatole degraders^15–19^. *Pseudomonas aeruginosa* Gs, isolated from mangrove sediment, can degrade up to 3 mM skatole within 8 days^17^. Additionally, two *Acinetobacter* species isolated from manure chickens were each capable of degrading 131 mg/L in 6 days^15^. In the strain *Cupriavidus* sp. KK10, skatole biotransformation occurs through different pathways, including carbocyclic aromatic ring fission, resulting in the formation of multiple products, such as single-ring pyrrole carboxylic acids^18^. Furthermore, *Burkholderia sp*. IDO3, isolated from activated sludge, was able to degrade 150 mg/L of skatole after 40 h^19^. RNA-seq analysis on the IDO3 strain revealed that skatole significantly regulated several oxidoreductase genes, including catechol 1,2-dioxygenase and catechol 2,3-dioxygenase. While many studies have identified microorganisms capable of utilizing skatole as a carbon source, our understanding of the specific genes involved in its degradation remains limited. Recently, the strain *Rhodococcus aetherivorans* DMU1, isolated from activated sludge, demonstrated the capability to utilize up to 25 mg/L of skatole as the sole carbon source^20^. Despite the elusive nature of the skatole metabolic pathway, recent findings indicate that an uncharacterized gene cluster in strain DMU1 may be involved in converting skatole to catechol^20,21^. This indicates a distinct and novel gene set dedicated to biodegradation, in contrast to those previously described for related compounds such as indole and indole-3-acetic acid (IAA)^22,23^.

We recently isolated a distinct species of *Rhodococcus*, *R. ruber* R1 originally identified for its ability to utilize methoxylated aromatic compounds as its sole carbon and energy source^24^. Here, we report the ability of *R. ruber* R1 to utilize skatole as its sole carbon source, underscoring its potential in biodegradation applications. We identify and characterize an operon of fourteen genes involved in the skatole degradation pathway. We describe aniline as a main intermediate in the route and six genes that encode an aniline dioxygenase system that converts aniline into catechol. The phylogenetic analysis shows that the aniline dioxygenase system is more broadly distributed among bacteria than previously thought, while the skatole catabolism pathway remains limited to certain species within the Actinomycetota phylum.

## RESULTS AND DISCUSSION

### Identification of the genetic cluster involved in skatole degradation in Rhodococcus ruber *R1*

*Rhodococcus ruber* R1, is a strain capable of utilizing various phenolic compounds as its sole carbon source for growth, including on the heterocyclic aromatic skatole (Fig. 1B). The strain was cultured in a minimal medium containing 1 mM skatole as the sole carbon and energy source for 30 hours. Skatole degradation was monitored through high-performance liquid chromatography (HPLC) analysis of the supernatants (Fig. 1B). This suggests that *Rhodococcus ruber* R1 contains a gene cluster specifically involved in the catabolism of skatole. To gain further insight into the skatole metabolic pathway, we performed a comparative genomic analysis of strain R1, contrasting it to other closely related *R. ruber* strains (Chol-4 and DSM43338) specifically chosen for their inability to grow on skatole (Fig. S1 and S2). This suggests that strain R1 may harbor skatole degradation genes not present in the other *R. ruber* species. Comparative analysis revealed that 420 genes are exclusively present in strain R1 (Fig. S1). Additionally, *R. aetherivorans* DMU1 was included in the analysis as it is capable of degrading skatole and carries an uncharacterized skatole degradation cluster^21^. Among the 420 genes unique to strain R1, 55 are shared with strain DMU1, including the set of previously identified genes in DMU1 potentially involved in skatole catabolism. These genes could also be implicated in skatole degradation in strain R1. This cluster consists of 14 genes, which we have designated as *skt* (Fig. S1). The operon comprises genes encoding a putative Rieske non-heme iron-dependent oxygenase (named *sktAB)* and a two-component flavoprotein monooxygenase (*sktGL)*. This is consistent with the presence of oxygenases in the catabolism of tryptophan-derived compounds, as previously described for indole and IAA^22,23^. The entire operon, encompassing 14 genes from *sktA* to *sktL* (Fig. S1), includes a hypothetical protein designated as *sktZ* and a potential transcriptional regulator named *sktR*.

To determine whether the *skt* genes identified in the genome of *R. ruber* R1 are implicated in skatole catabolism, we carried out a quantitative analysis by real-time PCR of RNA extracted from cells in the middle of the exponential phase. These cells were grown on skatole as the sole carbon and energy source, or fructose, as a non-related carbon source used as a control. Additionally, neighboring genes flanking the *skt* genes were analyzed, including two hypothetical proteins (E2561_RS07680 and E2561_RS07770), a putative pseudouridine-5’-phosphate glycosidase (E2561_RS07690) and a putative phosphodiesterase (E2561_RS07790). The results showed that the transcription levels of these neighboring genes did not change when exposed to skatole or fructose. In contrast, the transcription levels of the *sktA* gene (encoding the large subunit of the Rieske non-heme iron oxygenase) increased approximately 4000-fold in the presence of skatole (Fig. 2A). Similarly, the expression levels of *sktG and sktL* (encoding a putative two-component flavoprotein monooxygenase), and also *sktJ* (a hydantoinase-like enzyme) increased three orders of magnitude compared to the control (Fig. 2A). These results strongly suggest that the proteins from *sktA* to *sktL* genes may play a key role in the degradation of skatole (Fig. S1). Real-time PCR was also employed to analyze genes encoding catechol 1,2-dioxygenase, since catechol was proposed as an intermediate in skatole degradation in strain DMU1^20,21^.

**Figure 2.**
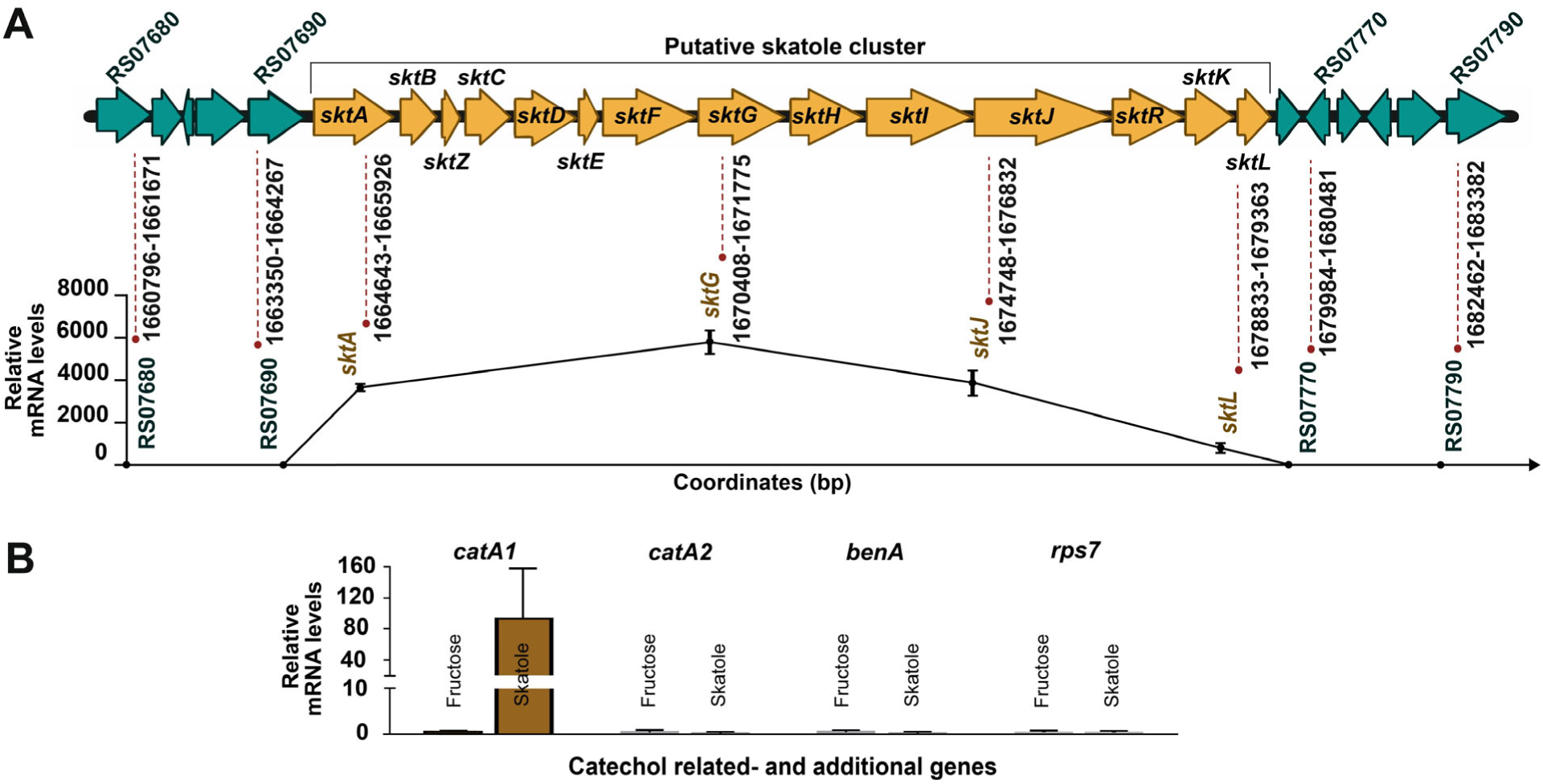
Transcript levels of skatole cluster measured in R1 strain exposed to skatole. **(A)** Real-time PCR analysis of *sktA*, *sktG*, *sktJ* and *sktL* genes expression from skatole cluster (yellow arrows, genes in bold), along with neighboring genes that flank the *skt* genes (green arrows, genes in bold). Cells were grown on skatole or fructose (control) as sole carbon source. Transcript levels were normalized to the average value observed in the fructose control treatment. 16S rRNA levels were used as an internal reference control. **(B)** Real-time PCR analysis of *catA1* (encoding putative catechol 1,2-dioxygenase), *catA2* (encoding a second putative catechol 1,2-dioxygenase), *benA* (encoding a large subunit of benzoate 1,2-dioxygenase), and *rpS7* (encoding ribosomal protein S7) genes. Cells were grown on skatole (brown bars) or fructose (control black bars) as sole carbon source. Transcript levels of *rpS7* and *benA* genes used as expression controls remained unchanged under these conditions. Three biological replicates were performed to determine the transcription levels. Error bars: ±SD.

As *R. ruber* R1 harbors two genes encoding catechol 1,2-dioxygenase, we examined both. Interestingly, only *catA1* (E2561_RS22180) showed increased expression in the presence of skatole compared to *catA2* (Fig. 2B), similar to observations described in DMU1^20,21^. Also, we used *benA* (encoding a large subunit of benzoate 1,2-dioxygenase), and *rpS7* (encoding ribosomal protein S7) as expression controls (Fig. 2B), which remained unchanged under the tested conditions.

It is worth noting that *R. aetherivorans* BCP1^25^, a bacterial *s*train similar to DMU1 that carries the *skt* cluster, was also capable of degrading skatole (Fig. S3). Collectively, the data strongly suggest that the *skt* genes are likely involved in skatole degradation not only in *Rhodococcus ruber* R1 but also in other *Rhodococcus* species. Likewise, these findings are consistent with a recent study on strain DMU1, where the *skt* cluster was identified through transcriptomic analysis^21^.

### Bioinformatic analysis of *skt* genes in *R. ruber* R1

To obtain more detailed information about *skt* genes (Fig. S1), we performed a comparison of the amino acid sequence of each protein encoded within the *skt* operon from R1 with those available in the UniProtKB/Swiss-Prot database. Results showed that both SktA and SktB exhibited the closest match (35% of amino acid identity) with the large subunit of 2-halobenzoate 1,2-dioxygenase from *Burkholderia cepacia* 2CBS^26^, and 35% of amino acid identity with the small subunit of p-cumate 2,3-dioxygenase from *Pseudomonas putida* F1^27,28^, respectively. These findings support the notion that SktAB encodes a Rieske non-heme iron-dependent oxygenase, comprising large and small subunits, similar to its detected homologous proteins. These types of proteins are part of a multi-component enzyme system that includes an electron transport chain and an oxygenase, catalyzing an array of oxidative transformations primarily related to the initial step in the degradation of aromatic compounds^29,30^. Intriguingly, the gene adjacent to *sktAB* is *sktZ* (Fig. S1), which is predicted to encode a hypothetical protein of 91 amino acids with no conserved domains, hence its function remains mysterious. In addition, results indicate that SktD *and* SktE encode a putative ferredoxin-NADP reductase and a ferredoxin, respectively. These proteins share 55% amino acid identity with a predicted ferredoxin-NADP reductase from *Frankia alni*^31^ and 66% amino acid identity with a putative ferredoxin from *Mycobacterium bovis*^32^. Both proteins seem to resemble an electron transport chain, which could be related to the oxygenase SktAB. Also, SktC exhibits low similarity (30% amino acid identity) with a glutamine amidotransferase (GAT) predicted automatically from *Methanopyrus kandleri*^33^, a function supported by analysis of conserved domains. This type of enzyme catalyzes the hydrolysis of glutamine, transferring the produced ammonia to a range of different metabolites^34^. Remarkably, SktF displays the highest similarity in the database (53% of amino acid identity) with a gamma-glutamylanide synthase (Glutamine synthetase-like protein (GS)) from *Acinetobacter sp*. YAA, which is part of an aniline oxidation cluster that catalyzes its conversion via gamma-glutamylanilide (γ-GA) into catechol^35,36^. Based on the previous information, collectively, *sktABCDEF* genes could encode a potential aniline dioxygenase. This assumption is drawn from the composition of this enzyme, which typically includes a GS-like protein, a GAT-like protein, an oxygenase, and a reductase^36,37^, characteristics shared with the aforementioned proteins in our strain. If this supposition holds validity, considering previous reports of aniline dioxygenase enzyme activity in Proteobacteria^37,38^, it is plausible that aniline and catechol may function as intermediates in the skatole pathway. This notion is further supported by findings in strain DMU1, which channeled skatole into catechol^20,21^, and induction of catechol dioxygenase in strain R1 in the presence of skatole (Fig. 2B).

Additional elements identified within the *skt* cluster include seven genes, designated *sktGHIJRKL* (Fig. S1). Briefly, SktG lacks a homolog in the UniProtKB/SwissProt database. Nevertheless, a putative conserved domain has been noticed, which is associated with the oxygenase component of styrene monooxygenase, a two-component flavoprotein monooxygenase enzyme^39,40^. This correlation is supported by the presence of a flavin reductase enzyme encoded by SktL, exhibiting 42% amino acid identity with the reductase component of p-hydroxyphenylacetate 3-hydroxylase from *Acinetobacter baumannii.* This flavoprotein contains FMN (flavin mononucleotide), and its function is to provide reduced flavin for the oxygenase component to hydroxylate its substrate^41,42^, thereby implying by analogy that SktGL could operate as a two-component flavoprotein monooxygenase. In addition, SktH appears to encode an enzyme belonging to the Xaa-Pro peptidase family, as deduced by conserved domain analysis, with no homologs found to be associated with this specific type of catabolism. Meanwhile, SktIJ displayed a similarity (34% and 42% amino acid identity) with the beta and alpha subunits of a caprolactamase from *Pseudomonas jessenii,* which is related to other ATP dependent enzymes involved in lactam hydrolysis, like 5 oxoprolinases and hydantoinases^43^. Besides, SktK contains a putative domain associated with short-chain dehydrogenases, a diverse family of oxidoreductases characterized by a single domain featuring a structurally conserved Rossmann fold and NAD(P)(H)-binding region. These enzymes catalyze various reactions, including isomerization, decarboxylation, epimerization, and carbonyl-alcohol oxidoreduction, among others ^44^. It is possible that these genes (*sktGHIJKL*) could encode enzymes responsible for catalyzing the conversion of skatole into aniline. Lastly, SktR, which encodes a putative LuxR regulator inferred through conserved domain analysis, completes the cluster (Fig. S1). All protein analyses were also supported by AlphaFold modeling (Fig. S4) and structure-based function prediction.

### Aniline as a potential intermediate in skatole degradation by *R. ruber* R1

To gather additional evidence regarding skatole degradation, we performed a more detailed examination of the growth of strain R1 on 1 mM skatole as the sole carbon or energy source. We collected samples at various time points and analyzed the supernatants using HPLC. As the biomass of *Rhodococcus ruber* R1 increased during growth on skatole, a corresponding decrease in skatole concentration was observed in the supernatants (Fig. 1B). Additionally, we observed the appearance and transient accumulation of an unknown compound (Fig. 1B, Fig. S5), which is of particular interest as it may represent a potential intermediate in the skatole degradation pathway. Based on previous bioinformatics analysis, aniline could potentially serve as an intermediate. To confirm this, we compared the retention time and UV spectrum of the unknown compound with an aniline standard using HPLC analysis. The results indicate that the unknown compound is indeed aniline (Fig. S5).

To determine if aniline was a substrate for R1 strain we performed an assay using R1 resting cells exposed to 1 mM aniline (Fig. 3A, black line). We did not observe a decrease in the aniline concentration after 6 h, suggesting that *R. ruber* R1 is either unable to degrade aniline or that this compound does not induce the cluster involved in its degradation. To determine if skatole was the cluster’s inducer, we performed a resting cell experiment pre-incubating *R. ruber* R1, with 0.5 mM skatole, following 1 mM aniline exposition (Fig. 3A, green line). We observed that after 6 h, the aniline concentration decreased by 100%, indicating that skatole induces aniline degradation in the strain *R. ruber* R1 (Fig. 3A).

**Figure 3.**
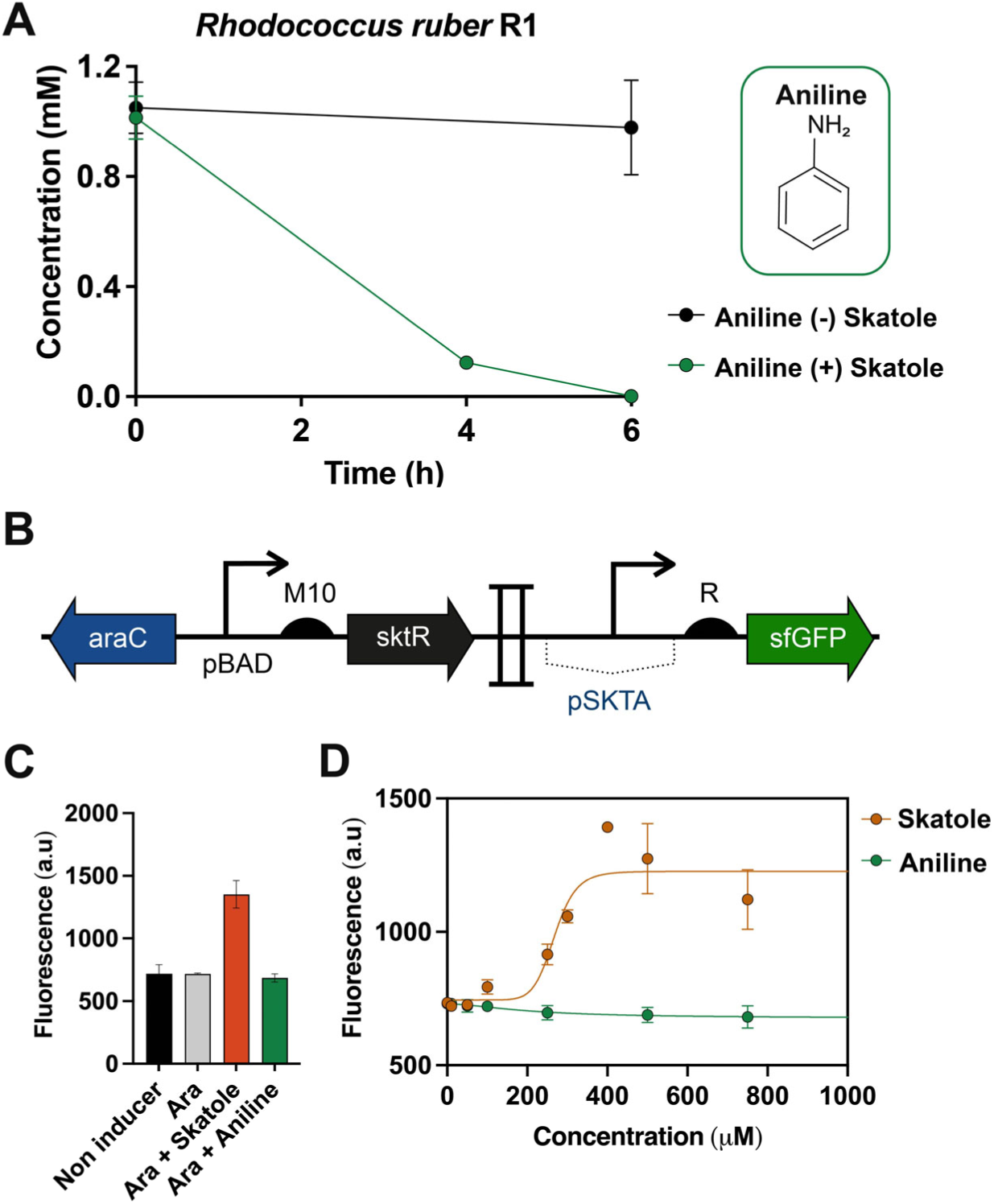
Specific induction of aniline degradation by the *skt* cluster in the presence of skatole. **(A)** Aniline degradation by *Rhodococcus ruber* R1 induced by skatole. Cells of *Rhodococcus ruber* R1 were grown on 20 mM fructose, induced with skatole (green line) or not (black line) then washed, and exposed to 1 mM aniline. High-performance liquid chromatography (HPLC) was performed to measure aniline degradation. Data points correspond to the mean value of the three replicates on three different resting cell experiments. **(B)** Response characterization of pSKTA promoter under skatole presence. Design of the heterologous reporter system in the strain *Cupriavidus pinatubonensis* JMP134. The reporter system was cloned in the inducible plasmid pBS1. SktR regulator was induced with 1 mM arabinose. **(C)** pSKTA promoter response to; non-inducer, arabinose 1mM, arabinose 1mM plus skatole 0.5 mM and arabinose 1 mM plus aniline 0.5 mM. Cells were grown in minimal medium A for 16 h at 30°C. Data points correspond to the mean value of the three replicates on three different days. Error bars: ±SD. **(D)** The response function of the reporter system to skatole or aniline. Cells were grown in minimal medium A with arabinose 1 mM and different skatole and aniline concentrations for 16 h at 30°C. The fit of the curve was obtained from the mean of three different experiments performed in triplicates on three different days. Error bars: ±SD.

To confirm that specifically, skatole induces its degradation cluster we built a heterologous reporter system in *Cupriavidus pinatubonensis* JMP134 by using the putative *skt* operon promoter (pSKTA) to control the expression of sfGFP, while including the putative SktR regulator under the control of pBAD promoter (Fig. 3B). Results showed that SktR activates the pSKTA promoter when induced with arabinose 1 mM and 0.5 mM skatole. However, no fluorescence induction was observed at 0.5 mM aniline (Fig. 3C). Next, we evaluated the dose-response curve of reporter strain JMP134 to skatole by flow cytometry. The reporter strain showed a clear skatole dose-dependent response with a detection limit of 50 µM (Fig. 3D). To determine the specificity of the SktR regulator toward skatole we also performed a dose-response curve to aniline, however, no response was observed, even at a high concentration such as 1 mM (Fig. 3D). These results confirm skatole as the main inducer of *skt* cluster and SktR as its regulatory protein.

As previously reported, aniline is converted into catechol through a series of steps by the aniline dioxygenase enzyme system in Proteobacteria^36,37^. Firstly, the GS-like protein (such as SktF) catalyzes the ATP-dependent ligation of L-glutamate to aniline, resulting in the formation of γ-GA. Subsequently, the oxygenase component (such as SktAB) accepts electrons from the reductase component (such as SktDE) to catalyze the conversion of γ-GA to catechol. Additionally, the GAT-like protein (such as SktC) can hydrolyze γ-GA to aniline and has been suggested to function as a detoxifying enzyme, as high concentrations of γ-GA are cytotoxic^36^. Notably, GS-like and GAT-like proteins are unique to this dioxygenase system and have not yet been identified in other aromatic compound dioxygenases.

To investigate whether SktABCDEF constitutes an aniline dioxygenase system, we initially cloned the *sktF* gene into a heterologous strain and analyzed the conversion of aniline into γ-GA. Our results showed that heterologous cells expressing the *sktF* gene, when exposed to 1 mM aniline and 1 mM glutamate (Fig. S6A, left panel), exhibited a reduction in aniline concentration of approximately 30% after 6 hours, accompanied by the formation of γ-GA. In contrast, cells exposed to 1 mM aniline alone (Fig. S6A, right panel) showed no detectable formation of γ-GA after 6 hours. These findings strongly suggest that SktF may act as a γ-GA synthetase, which also requires the presence of glutamate for its function, similar to previously described γ-GA synthetases in Proteobacteria^37^. Altogether these data confirm aniline as an intermediate in the degradation pathway of skatole.

### Heterologous expression and resting cell assays suggest a key role of *sktABCDEF* from *R. ruber* R1 in the degradation of aniline and its derivatives

To validate the previously assumed function of SktABCDEF as an aniline dioxygenase system capable of converting aniline into catechol (Fig. 4A), we cloned and expressed these genes in the heterologous strain *C. pinatubonensis* JMP134. The results demonstrated that the heterologous strain containing the *sktABCDEF* genes (designated as pBS1-sktA-F) and cultured on aniline supplemented with an additional carbon source (fructose) exhibited a dark coloration (on the right side of the plate) (Fig. 4B). This coloration is indicative of catechol accumulation from aniline, followed by its subsequent polymerization. This observation is supported by previous findings suggesting that catechol polymerizes upon exposure to oxygen, resulting in the formation of a dark brown pigment^45^. In contrast, the heterologous strain carrying an empty vector (on the left side of the plate) did not manifest any dark coloration (Fig. 4B). In addition, the heterologous strain harboring pBS1-sktA-F (green line), incubated in liquid cultures containing aniline as the sole nitrogen source supplemented with fructose as the sole carbon source, exhibited a significant growth (Fig. 4C). On the other hand, the strain carrying the empty vector failed to proliferate (black line) (Fig. 4C). These findings imply that the conversion of aniline into catechol occurs concurrently with the release of ammonia in the strain harboring the *sktABCDEF* genes, as previously documented for aniline dioxygenases^36,46^. Taken together, these findings confirm aniline as an intermediate in the skatole degradation pathway and underscore the critical role of the *sktABCDEF* cluster in encoding an aniline dioxygenase (Fig. 5A).

**Figure 4.**
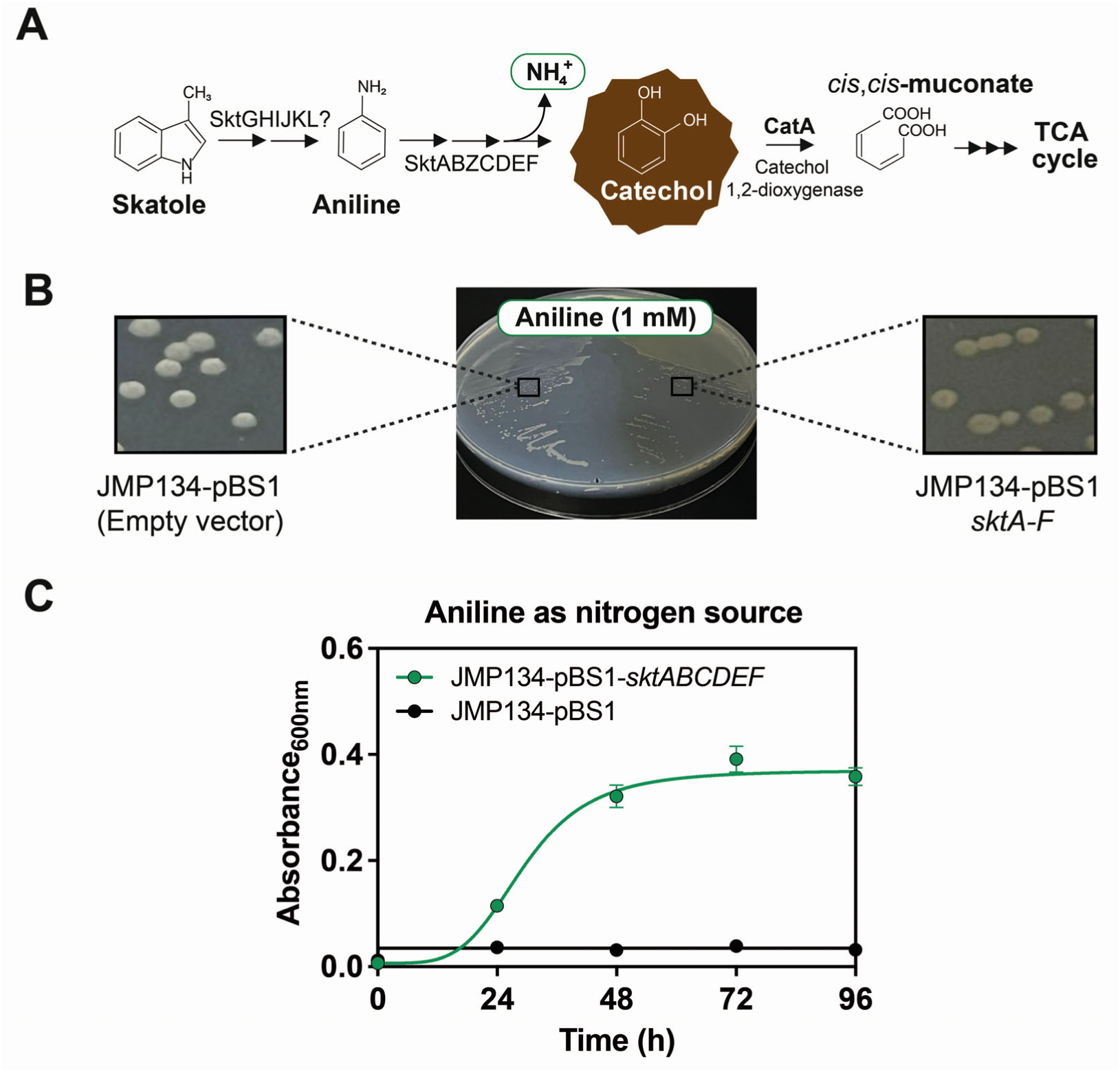
Expressing putative aniline dioxygenase genes (*sktA-F*) in the heterologous strain JMP134. **(A)** The putative *R. ruber* R1 route channeling degradation of skatole through aniline and catechol is illustrated. This reaction release ammonium. Catechol is depicted surrounded by a brown image because it is known to polymerize upon contact with O_2_, producing a brown dark pigment. **(B)** Growth of heterologous strain JMP134 harboring *sktABZCDEF* genes (on the right side, JMP134-pBS1-*sktA-F*) or JMP134 carrying the empty vector (on the left side, JMP134-pBS1) containing aniline plus an additional carbon source (fructose). A dark coloration in the strain harboring putative aniline dioxygenase genes is noted. **(C)** Growth on a liquid culture of JMP134-pBS1-*sktA-F* (green line) on 1 mM aniline as sole nitrogen source compared to empty vector (black line). 10 mM fructose was used as a carbon source. Three biological replicates were performed for growth measurements. Error bars: ±SD.

**Figure 5.**
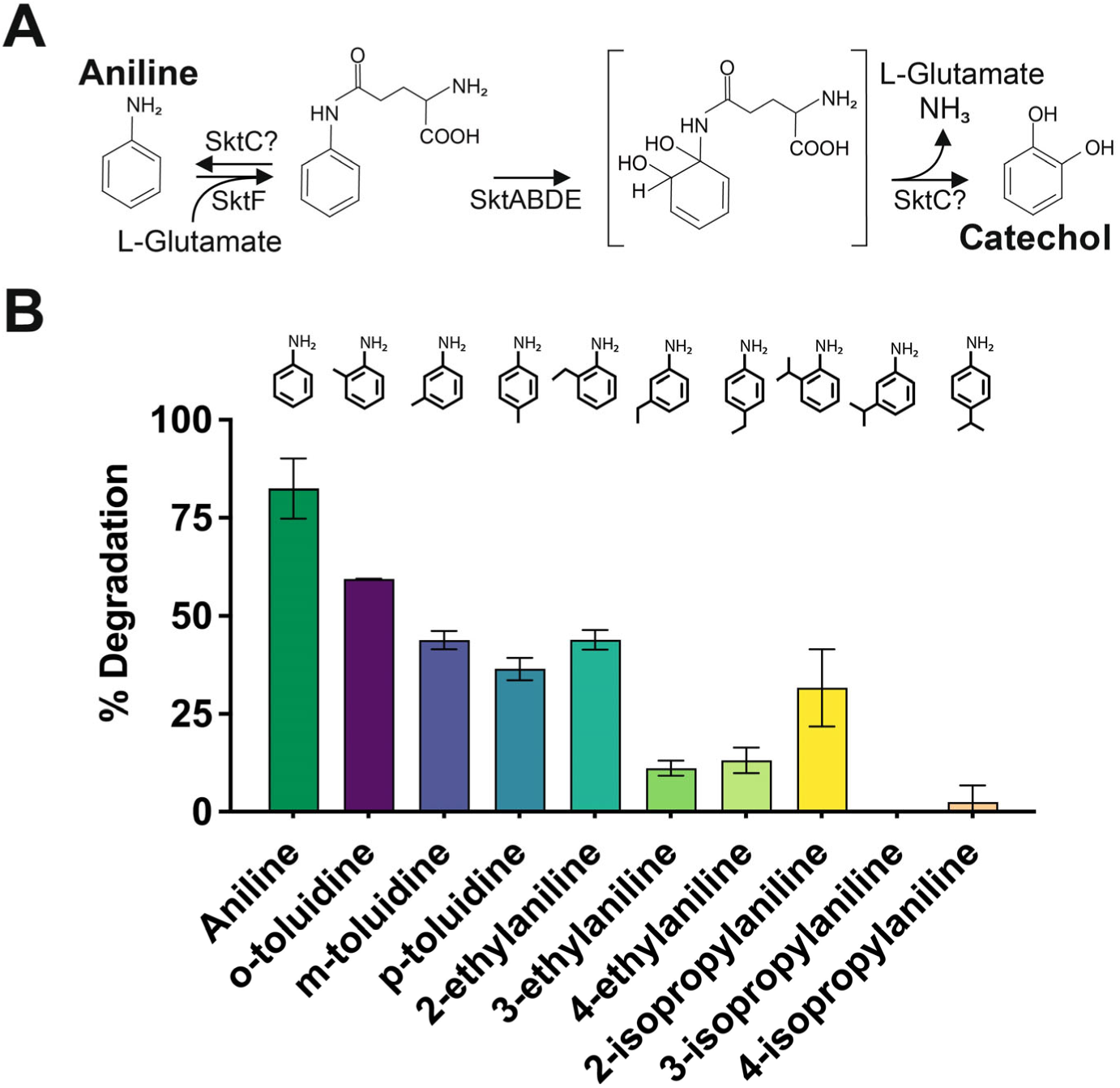
Role of the *sktABCDEF* cluster encoding an aniline dioxygenase in degradation of aniline and its derivatives. **(A)** Proposed pathway for the degradation of aniline in the Actinobacterium *Rhodococcus ruber* R1. SktF and SktC are GS-like and GAT-like proteins, respectively, while SktABDE constitutes the dioxygenase system. This pathway has been adapted from previous reports in Proteobacteria. **(B)** Degradation percentages of aniline and derivatives (toluidines, ethylanilines, and isopropylanilines) by *C. pinatubonensis* JMP134 expressing *sktA-F* genes. Cells of *C. pinatubonensis* JMP134 harboring the inducible cluster *sktABZCDEF* were grown on 20 mM fructose supplemented with 2.5 mM arabinose as an inducer for 3 h. The cells were then washed and subsequently exposed to 1 mM aniline and derivatives for 2h. Three biological replicates were performed to determine the percentage degradation. Error bars: ±SD.

Earlier investigations have demonstrated the conversion of aniline using an extracted cell of *E. coli* SO58 containing AtdA1 protein (similar to that encoded by *sktF*), achieving an 80% reduction in aniline within 8 hours. They also showed reductions in its methylated derivatives, *o*-methylaniline (40%), *m*-methylaniline (27%), and *p*-methylaniline (45%)^36^. Based on this, we tested heterologous cell suspensions expressing *sktA-F* genes. In this case, our cells were induced with arabinose for 3 hours using a pBAD promoter to regulate the expression of the entire aniline dioxygenase system. After induction, the cells were washed and then exposed to 1 mM aniline and its derivatives for 2 hours (Fig. 5B). These results showed that aniline was almost completely degraded (82%), whereas substituted methylanilines at the *o-*, *m-*, and *p-* positions were degraded by 59%, 44%, and 36%, respectively (Fig. 5B). These proportions were relatively close to those reported in Takeo *et al*^36^. In addition to methylanilines, other anilines such as ethylanilines and isopropylanilines at the *o-, m-*, and *p-* positions were also analyzed. 2-ethylaniline was degraded by 43%, whereas 3-ethylaniline and 4-ethylaniline were degraded by 11% and 13%, respectively. 2-isopropylaniline was degraded by 32%, 4-isopropylaniline was degraded by only 2%, and 3-isopropylaniline was not degraded. These data suggest that the aniline dioxygenase enzyme system is more permissive with less bulky substituents, and it also shows a preference for the *o*-position, indicating that *o*-substituted anilines also function as substrates, albeit with lower affinity.

The provided data confirm that the *sktABCDEF* genes of *R. ruber* R1 encode an aniline dioxygenase system, which can catalyze the conversion of aniline and its derivatives.

### Unraveling the phylogeny of the aniline dioxygenase system in *Rhodococcus ruber* R1

As previously reported by Ji *et al*^37^, the substrate specificity of aniline dioxygenases from *Sphingobium* and *Acinetobacter* primarily relies on two key components: the glutamine synthetase-like enzyme and the oxygenase. These components also exhibit a more conserved amino acid composition among these bacteria compared to the other subunits^37^, likely due to their direct interaction with substrates. This suggests that this applies to all aniline dioxygenases that share a similar composition. Therefore, to assess the presence of aniline dioxygenase genes in bacterial genomes, we selected SktA as the gene marker, corresponding to the large subunit of the oxygenase component of aniline dioxygenase from strain R1. This choice is based on its expected essential role in substrate recognition and catalysis, as well as the degree of amino acid conservation.

Then, a search for SktA homologs in available bacterial genomes revealed the presence of this protein in diverse bacterial species spanning the Actinomycetota, Pseudomonadota, Bacillota, and Cyanobacteriota phyla isolated from diverse sources (Fig. 6, Fig. S7). The number of phyla carrying SktA homologs indicates a broader distribution of this aniline dioxygenase system among bacteria than initially thought^47^. A more in-depth analysis of the strains harboring the putative large subunit of the oxygenase component of aniline dioxygenase, revealed that in the majority of cases, the small subunit of this oxygenase, along with the γ-GA synthetase and a GAT-like protein, are located in close proximity to the analyzed gene (*sktA*-like) (Fig. S7).

**Figure 6.**
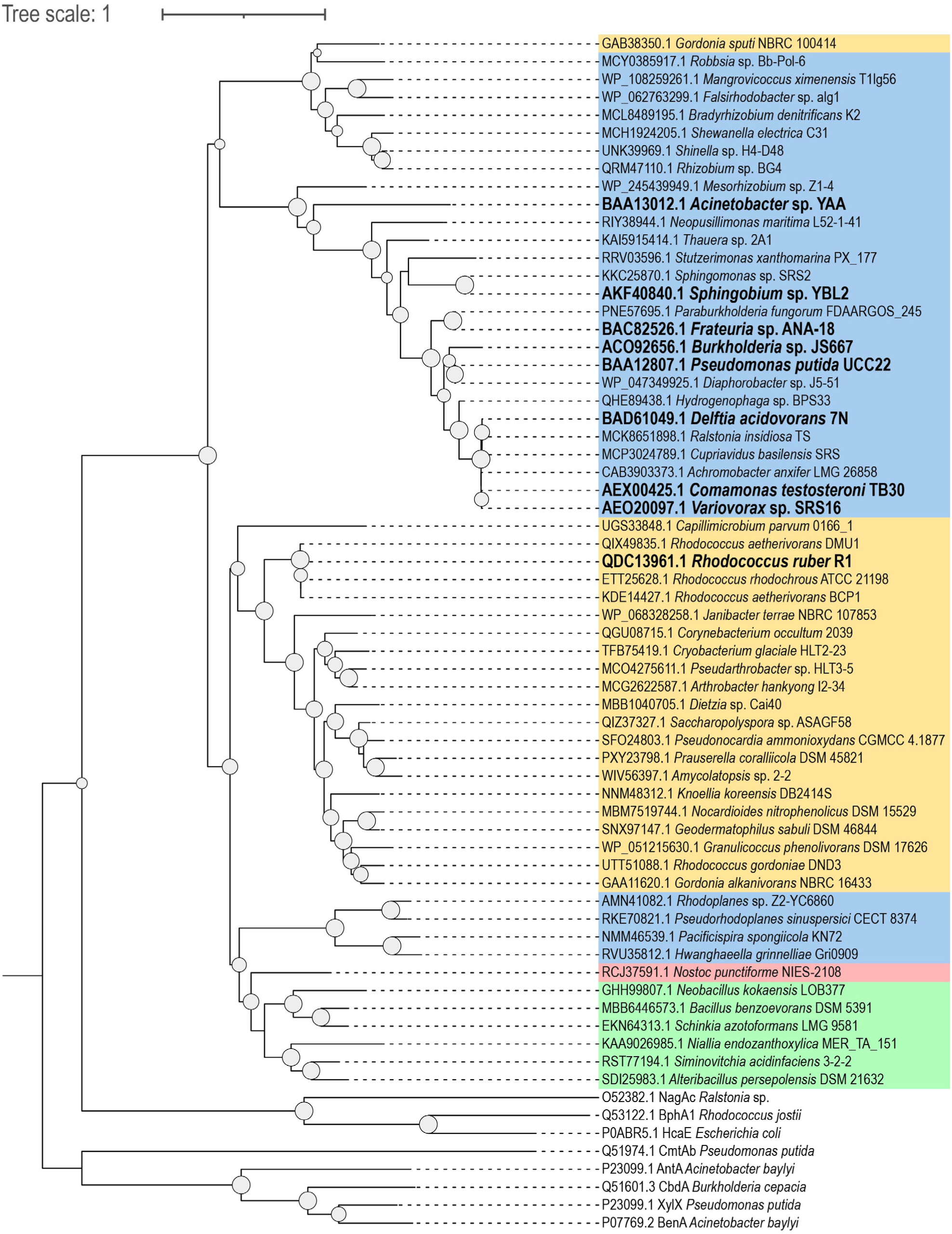
Evolutionary relationships among large subunit of aniline dioxygenase (SktA) homologs from selected bacterial species. Putative large subunit of aniline dioxygenase proteins were analyzed using the Maximum Likelihood topology provided by IQ-TREE software. Sequence alignments were computed using MAFFT software, and support values for likelihood ratio tests similar to SH (n=1000) were assigned to each node (values >50% are shown). For the outgroups, oxygenase components from various organisms were utilized, including 3-phenylpropionate/cinnamic acid dioxygenase (*Escherichia coli* K12), biphenyl 2,3-dioxygenase (*Rhodococcus jostii* RHA1), naphthalene 1,2-dioxygenase (*Ralstonia* sp. U2), benzoate 1,2-dioxygenase (*Acinetobacter baylyi* ADP1), anthranilate 1,2-dioxygenase (*A. baylyi* ADP1), toluene 1,2-dioxygenase (*Pseudomonas putida*), and 2-halobenzoate 1,2-dioxygenase (*Burkholderia cepacia* 2CBS). Bold-highlighted sequences indicate strains known to possess aniline dioxygenase activity, either demonstrated biochemically or genetically. Taxonomic classification is represented by a color scheme: light blue for Pseudomonadota, yellow for Actinomycetota, green for Bacillota, and light red for Cyanobacteriota.

It is interesting to note that the electron transport chain supporting the function of oxygenase appears to vary among different phyla. In the predominant number of investigated Pseudomonadota, we identified a single iron-sulfur flavoprotein, analogous to phthalate dioxygenase reductase^48^, in the genomic context of the aniline dioxygenase gene, consistent with previous descriptions^46,47,49–51^. However, in Actinomycetota, we identified near the *sktA*-like gene an electron transport chain consisting of two components: a flavoprotein (ferredoxin-NADP reductase) and an iron-sulfur protein [2Fe–2S] (ferredoxin-type protein). These components are similar to Rhodocoxin reductase-Rhodocoxin partners found in *Rhodococcus erythropolis*, which are involved in transferring electrons from NADH to cytochrome P450 during the degradation of thiocarbamate herbicides^52^. A notable case is found in strain *Capillimicrobium parvum* 0166_1^53^, which harbors a ferredoxin with a 4Fe-4S single cluster domain, as deduced from conserved domain analysis. Interestingly, it is worth noting that only *Rhodococcus* species exceptionally possesses an NAD(P)/FAD-dependent oxidoreductase similar to Thioredoxin reductases, as well as a putative ferredoxin containing a novel 4Fe-4S dicluster domain, inferred by conserved domain analysis. This finding is in concordance with plant-type ferredoxin:thioredoxin reductase and methanogenic archaeal ferredoxin:disulfide reductase, both of which feature an unusual active-site [4Fe-4S] cluster^54–56^. These data indicate that Actinomycetota possess a six-component aniline dioxygenase system, differing from that of the Pseudomonadota phylum. *Rhodococcus* species, in particular, represent a special case for future studies related to the unique reductase subunit. Regarding the Bacillota and Cyanobacteriota phyla, no evident electron transport chain was found in close proximity to putative aniline dioxygenase systems (Fig. S7), and its composition remains elusive.

Furthermore, the hypothetical protein encoded by *sktZ* was found in Actinomycetota and Bacillota, but not in Pseudomonadota or Cyanobacteriota (Fig. S6). AlphaFold modeling (Fig. S4) and structure-based function prediction^57^ for this protein provided insights into its function, which has been classified as a cellular component with a CscoreGO (GO terms) of 0.59^57^. In conjunction with the observation that this protein was identified only in Actinomycetota and Bacillota, SktZ could be functionally related to a structural role in Gram-positive bacteria.

Remarkably, among the selected bacteria harboring the aniline dioxygenase system, only *Cryobacterium glaciale* HLT2-23^58^, *Arthrobacter hankyongi* I2-34^59^ and *Pseudarthrobacter sp*. HLT3-5 (GCA_023913815.1) carries additional *skt* genes (*sktGHIJKRL*) (Fig. S7). This suggests that other bacteria possessing the aniline dioxygenase system may be exclusively dedicated to metabolizing this substrate or channeling another compound towards aniline or their derivatives, while the capability to degrade skatole could be limited to specific genera from Actinomycetota phylum. To confirm that aniline is a substrate for bacteria carrying incomplete *skt* cluster, we analyzed the ability to degrade this substrate using *Gordonia alkanivorans* NBRC 16433^60^, a bacterium belonging to the Actinomycetota phylum, which only carries the aniline degradation genes (Fig. S7). This bacterium effectively degrades aniline completely in a resting cell assay (Fig. S8). These findings suggest that the identified cluster constitutes a genuine aniline dioxygenase system within this phylum. This conclusion further underscores the ubiquitous presence of anilines in the environment and suggests their potential significance as natural substrates. Additionally, anilines and their derivatives are widely used in the production of dyes, pharmaceuticals, and agricultural chemicals. Their release into the environment, primarily through industrial effluents and agricultural runoff, poses significant ecological and health risks^61,62^. Certain anilines, such as *o-*, *m-*, and *p-*toluidine, are recognized as carcinogenic, mutagenic, and teratogenic, and are listed as priority pollutants by the United States Environmental Protection Agency (EPA)^63,64^. The removal of anilines from the environment is essential, and this study provides insights into the genes involved in aniline degradation in Actinomycetota. This information facilitates the development of bioremediation strategies for environments contaminated with aniline and skatole and advances the design of skatole biosensors through the targeted manipulation of the identified promoter and regulatory protein associated with the *skt* cluster.

## 4. CONCLUSIONS

In summary, we have confirmed the existence of a new genetic cluster in *Rhodococcus ruber* R1 crucial for skatole degradation, composed of 14 genes (Fig. S1). Among them, the *sktABCDEF* genes encode an aniline dioxygenase enzyme system analogous to those previously described in Pseudomonadota. Through resting cellular metabolism assays and subsequent HPLC-UV analysis of the supernatants, we verified that this enzyme system effectively degrades aniline and methylanilines. Additionally, we demonstrate that aniline serves as an intermediate in skatole metabolism, and its enzyme system that converts aniline into catechol plays a crucial role in this catabolic process. Further, we identified the promoter and regulatory protein associated with the *skt* genes, which are induced by skatole. Phylogenetic analysis indicates that skatole degradation-related genes appear to be restricted to certain species within the Actinomycetota phylum, whereas aniline biodegradation could be more widespread across bacteria than previously thought.

## MATERIALS AND METHODS

### Bacterial strains, plasmids, and growth conditions

Bacterial strains and plasmids used in this study are detailed in Table S1. *Rhodococcus ruber* R1, *R. ruber* chol-4, *R. ruber* DSM 43338, *R. aetherinovorans* BCP1, *Gordonia alkanivorans* NBRC 16433, *Cupriavidus pinatubonensis* JMP134 and *C. pinatubonensis* JMP222 were cultured at 30°C in mineral salt medium (MSM) as previously described by Donoso et al 2021^65^, supplemented with 1 mM skatole, 1 mM aniline, or 5 mM fructose as the sole carbon and energy sources. Derivative strains of *C. pinatubonensis* JMP134 were cultivated at 30 °C in either MSM or in nitrogen-free MSM (modified from Donoso *et al.,* 2021^65^, substituting Ca(NO_3_)_2_•4H_2_O and (NH_4_)_2_SO_4_ with CaSO_4_) supplemented with 1 mM aniline as the sole nitrogen source, 20 mM fructose as the main carbon source, and gentamicin (20 µg/ml) when appropriate. Growth was monitored by absorbance at 600 nm (OD600) using a Spectroquant® Prove spectrophotometer (Merck, Darmstadt, Germany). Each growth measurement was performed with at least three biological replicates.

### Detection of transcripts by quantitative real-time PCR

Cells of *R. ruber* R1 were cultured using 1 mM skatole or 5 mM fructose as the sole carbon and energy sources. Total RNA was extracted from 4 mL of mid-log-phase cells using Trizol (Invitrogen, Carlsbad, CA). The RNA was quantified with an Eon microplate spectrophotometer (BioTek, Winooski, VT, USA) and treated with the Turbo DNase kit (Thermo Fisher, Massachusetts, USA) to remove DNA contamination. Reverse transcription PCR was conducted with the ImProm-II reverse transcription system (Promega Corporation, Madison, WI, USA) using 1 µg of RNA in 20 µL reaction mixtures. Real-time PCR was performed using the Brilliant II SYBR Green qPCR Master Mix (Agilent Technologies). The PCR mixture (15 µL) contained 3 µL of template cDNA (diluted 1:10) and 0.2 µM (each) primer. Amplification conditions were 95°C for 5 min, followed by 40 cycles of 95°C for 30 s, 60°C for 30 s, and 72°C for 40 s, with a final melting cycle from 55°C to 95°C. Relative gene expression values were calculated using the comparative cycle threshold method (2^-ΔΔCT^ method)^66^, with the 16S rRNA (locus_tag: E2561_RS01515, E2561_RS02600, E2561_RS08625, E2561_RS12595) gene sequence used as a reference gene (internal control). Gene expression levels were normalized to the average value of the gene expression levels determined in the fructose treatment. Genes analyzed by quantitative real-time PCR included RS07680 (locus_tag: E2561_RS07680), RS07690 (E2561_RS07690), *sktA* (E2561_RS07695), *sktG* (E2561_RS07730), *sktJ* (E2561_RS07745), *sktL* (E2561_RS07760), RS07770 (E2561_RS07770), RS07790 (E2561_RS07790), *catA1* (E2561_RS22180), *catA2* (E2561_RS18270), *benA* (E2561_RS22230) and *rps7* (E2561_RS16860). Primer pairs used are listed in Table S2. Experiments were performed in triplicate.

### Plasmids construction

A plasmid containing the entire region between *sktA* and *sktF* genes, regulated by an L-arabinose-inducible promoter, designated as pBS1-*sktA-F* (Table S1), was constructed using the Gibson assembly method^67^. Briefly, PCR products comprising *sktABZCDEF* genes and pBS1 plasmid were generated using primers pairs sktA-Fw1/sktCRv1, sktC-Fw2/sktF-Rv2 and pBS1a/pBS1b (listed in Table S2). These primers contain a 20-bp terminal sequence homologous to the terminus of the fragment to be linked. The sequences were combined and ligated to generate a new DNA molecule (pBS1-*sktA-F*) in a one-step isothermal reaction^67^. Plasmid pBS1 derivatives were transformed into *E. coli* Mach1, and the transformed cells were selected in LB medium supplemented with gentamicin (20 µg/mL). Proper insertion of the *sktA-F* genes was verified by PCR, and the full-length gene construct was confirmed by Sanger sequencing to ensure the absence of errors. The same procedure was utilized to construct a plasmid containing only the *sktF* gene (Table S1), using the primer pair sktF-Fw3/sktF-Rv3 (Table S2) in combination with primers for pBS1 amplification (Table S2). Subsequently, the pBS1-derived plasmids were transformed into the JMP134 or JMP222 strains for phenotypic analysis.

### Promoter characterization

The reporter device was built on the backbone pBS1 containing a pBRR1 origin of replication and a gentamicin resistance cassette. To construct the reporter device the sequence of the putative wild-type promoter of the *sktA* gene was synthesized by IDT Technologies, together with an *E. coli*, codon-optimized version of *sktR* gene. All primers were designed to support cloning by Gibson assembly^67^. DNA sequences are provided in the Supporting Information. For cloning, DH5alphaZ1 was grown on LB medium supplemented with 30 μg/mL gentamicin. The plasmid was purified using the QIAprep spin Miniprep kit (Qiagen) and sequence-verified by Sanger sequencing (Eurofins Genomics, EU). To perform the promoter induction measurements, three different colonies of the JMP134 strain harboring the reporter device were picked and inoculated, separately, into 500 μL MSM supplemented with 10 mM fructose and 30 μg/mL gentamicin in 96 DeepWell polystyrene plates (Thermo Fisher Scientific, 278606) sealed with an AeraSeal film (Sigma-Aldrich, A9224-50EA) and incubated at 30°C for 16h with shaking and 80% humidity in a Kuhner LT-X (Lab-Therm) incubator shaker. After overnight growth, the cells were diluted 100 times into a fresh MSM with antibiotics, 10 mM fructose, and skatole or aniline at different concentrations. The cells were induced at 30 °C for 16 h with shaking. The cells were diluted 100 times in PBS and analyzed by flow cytometry. All experiments were performed in triplicate on three independent days.

### Resting cell assays and analytical methods

To determine the concentration of aniline, 2-methylaniline (*o*-toluidine), 3-methylaniline (*m*-toluidine), 4-methylaniline (*p*-toluidine), 2-ethylaniline, 3-ethylaniline, 4-ethylaniline, 2-isopropylaniline, 3-isopropylaniline, 4-isopropylaniline, and L-glutamic acid gamma-anilide, cell-free supernatants from resting cells of *Rhodococcus ruber* R1 grown on 20 mM fructose overnight, JMP134, JMP222, and its derivatives grown on 20 mM fructose overnight and induced for 3 h with 2.5 mM arabinose, were analyzed by high-performance liquid chromatography (HPLC), following the method described by^68^. Briefly, after washing JMP134 cells and their derivatives twice with 1 volume of MSM, they were concentrated 10 times and incubated with 1 mM of each compound mentioned above. JMP222 cells and their derivatives were used at 1-fold concentration and incubated with L-glutamic acid gamma-anilide, and glutamate if appropriate. Samples (200 μl) were taken at different times, filtered (0.22 μm), and injected into a JASCO LC-4000 liquid chromatography (JASCO, Oklahoma City, OK, USA) equipped with a Kromasil 100-3.5-C18 column (4.6 mm diameter). Chromatographic conditions employed a mobile phase of methanol/water (50% v/v) at a flow rate of 0.8 ml/min. The effluent from the column was monitored at 232 nm wavelengths for the detection of whole compounds, including aniline (t_R_ = 3.5 min), *o*-toluidine (t_R_ = 5.2 min), *m*-toluidine (t_R_ = 5.2 min), *p*-toluidine (t_R_ = 5.3 min), 2-ethylaniline (t_R_ = 8.5 min), 3-ethylaniline (t_R_ = 8.9 min), 4-ethylaniline (t_R_ = 9.4 min), 2-isopropylaniline (t_R_ = 13.7 min), 3-isopropylaniline (t_R_ = 15.2 min), 4-isopropylaniline (t_R_ = 15.7 min) and L-glutamic acid gamma-anilide (t_R_ = 2.2 min).

### Bioinformatic tools

To analyze and characterize the skatole-degrading genes in *Rhodococcus* species, we have used several bioinformatic tools and techniques:

#### Genome Annotation and Pan-genome Analysis

The genomic sequences of *Rhodococcus* species in FASTA format were re-annotated using Prokka v1.14.6 with default settings^69^. Roary v3.11.2 was employed to classify the obtained genes as core, cloud, or shell with a gene classification threshold of 90% identity^70^. Default parameters were used for the remaining settings.

#### Gene Sequence Retrieval

SktA homologous genes were retrieved from the most recent version of the GenBank non-redundant protein sequence database^71^. Proteins that had a minimum of 45% amino acid identity to SktA of *Rhodococcus ruber* R1 were taken into thorough consideration.

#### Phylogenetic Analysis

The sequences were aligned using MAFFT v7.475 under default settings, the best practice option for very accurate alignment of amino acid sequences^72^. A maximum likelihood phylogenetic tree was done by the software IQ-TREE v1.6.12. Support values were obtained using the SH-like method of 1000 replicates of bootstrap in the likelihood ratio tests to assign confidence levels to the nodes of the tree^73^. Visualization and editing of phylogenetic trees were performed by the Interactive Tree Of Life (iTOL) online tool^74^.

#### Functional Annotation

The function of the identified proteins was predicted concerning conserved domains using the UniProtKB/Swiss-Prot database. The *skt* genes obtained from *Rhodococcus ruber* R1 were examined through this database to detect homologous proteins, from which their functions were inferred. Identification of conserved domains within a protein was performed using the Conserved Domains Database from the NCBI website^75^. Additionally, we employed AlphaFold^76^ with default parameters as implemented in AlphaFold2.ipynb available on ColabFold v1.5.5 to predict protein structures, and we used COFACTOR^57^ to infer their functions.

#### Genomic Context Analysis

Genomic context and organization of the *skt* genes were determined to identify functional neighboring genes and operons. Comparative genomics was performed on the R1 strain (ASM635198v2) and other *Rhodococcus* strains, both degrading (DMU1, ASM1227295v1) and non-degrading skatole (Chol-4, ASM34795v2; DSM43338, ASM164683v1).

### Chemicals

Skatole, Aniline, 2-Methylaniline, 3-Methylaniline, 4-Methylaniline, 2-Ethylaniline, 3-Ethylaniline, 4-Ethylaniline, 2-Isopropylaniline, 3-Isopropylaniline, 4-Isopropylaniline and L-Glutamic acid were purchased from Sigma-Aldrich (Steinheim, Germany). L(+)-Arabinose, Benzoic acid and Fructose were purchased from Merck (Darmstadt, Germany). L-Glutamic acid gamma-anilide was purchased from Santa Cruz Biotechnology (Santa Cruz, CA, USA).

## Supporting information

Supplementary information, Galaz et al 2024

## AUTHOR CONTRIBUTIONS

R.D. designed the research. S.G.J. conducted all experiments related to qPCR, gene cloning, plasmid construction, resting cell assays, and analytical methods. B.S. provided support for the aforementioned techniques. F.G.T. performed bioinformatic analysis. A.Z., carried out experiments related to promoter characterization. S.G.J., R.D and A.Z. analyzed the data, and wrote the manuscript. R.D. and A.Z. edited the article. All authors reviewed and approved the manuscript.

## DATA AVAILABILITY STATEMENT

The data supporting this work’s conclusions are included within the manuscript, and no large datasets were generated or analyzed during the current study.

## ACKNOWLEDGMENTS

Some figures were created using BioRender.com.

## REFERENCES

1. Claus, R., Weiler, U. & Herzog, A. Physiological aspects of androstenone and skatole formation in the boar-A review with experimental data. Meat Sci. 38, 289–305 (1994).

2. Deslandes, B., Gariépy, C. & Houde, A. Review of microbiological and biochemical effects of skatole on animal production. Livest. Prod. Sci. 71, 193–200 (2001).

3. Liu, D. et al. Indoleacetate decarboxylase is a glycyl radical enzyme catalysing the formation of malodorant skatole. Nat. Commun. 9, 4224 (2018).

4. Duca, D., Lorv, J., Patten, C. L., Rose, D. & Glick, B. R. Indole-3-acetic acid in plant-microbe interactions. Antonie Van Leeuwenhoek 106, 85–125 (2014).

5. Tang, J. et al. Biosynthetic pathways and functions of indole-3-acetic acid in microorganisms. Microorganisms 11, (2023).

6. Cohen, J. D. & Strader, L. C. An auxin research odyssey: 1989-2023. Plant Cell 36, 1410–1428 (2024).

7. Attwood, G., Li, D., Pacheco, D. & Tavendale, M. Production of indolic compounds by rumen bacteria isolated from grazing ruminants. J. Appl. Microbiol. 100, 1261–1271 (2006).

8. Lee, M.-H. et al. Effect of slurry treatment approaches on the reduction of major odorant emissions at a hog barn facility in South Korea. Environ. Technol. 38, 506–516 (2017).

9. Trabue, S., Kerr, B., Bearson, B. & Ziemer, C. Swine odor analyzed by odor panels and chemical techniques. J. Environ. Qual. 40, 1510–1520 (2011).

10. Zhou, Y., Hallis, S. A., Vitko, T. & Suffet, I. H. M. Identification, quantification and treatment of fecal odors released into the air at two wastewater treatment plants. J. Environ. Manage. 180, 257–263 (2016).

11. Simeoni, M. et al. Correction to: An open-label, randomized, placebo-controlled study on the effectiveness of a novel probiotics administration protocol (ProbiotiCKD) in patients with mild renal insufficiency (stage 3a of CKD). Eur. J. Nutr. 58, 2157 (2019).

12. Lombardi, F. et al. The effects of low-nickel diet combined with oral administration of selected probiotics on patients with systemic nickel allergy syndrome (SNAS) and gut dysbiosis. Nutrients 12, 1040 (2020).

13. Hammouda, S. B., Adhoum, N. & Monser, L. Synthesis of magnetic alginate beads based on Fe3O4 nanoparticles for the removal of 3-methylindole from aqueous solution using Fenton process. J. Hazard. Mater. 294, 128–136 (2015).

14. Fischer, J. et al. Fast and solvent-free quantitation of boar taint odorants in pig fat by stable isotope dilution analysis-dynamic headspace-thermal desorption-gas chromatography/time-of-flight mass spectrometry. Food Chem. 158, 345–350 (2014).

15. Tesso, T. A., Zheng, A., Cai, H. & Liu, G. Isolation and characterization of two *Acinetobacter* species able to degrade 3-methylindole. PLoS One 14, e0211275 (2019).

16. Sharma, N., Doerner, K. C., Alok, P. C. & Choudhary, M. Skatole remediation potential of *Rhodopseudomonas palustris* WKU-KDNS3 isolated from an animal waste lagoon. Lett. Appl. Microbiol. 60, 298–306 (2015).

17. Yin, B. & Gu, J.-D. Aerobic degradation of 3-methylindole by *Pseudomonas aeruginosa* Gs isolated from mangrove sediment. Hum. Ecol. Risk Assess. 12, 248–258 (2006).

18. Fukuoka, K., Ozeki, Y. & Kanaly, R. A. Aerobic biotransformation of 3-methylindole to ring cleavage products by *Cupriavidus* sp. strain KK10. Biodegradation 26, 359–373 (2015).

19. Ma, Q. et al. Biodegradation of skatole by *Burkholderia* sp. IDO3 and its successful bioaugmentation in activated sludge systems. Environ. Res. 182, 109123 (2020).

20. Ma, Q. et al. Unraveling the skatole biodegradation process in an enrichment consortium using integrated omics and culture-dependent strategies. J. Environ. Sci. (China) 127, 688–699 (2023).

21. Li, Y. et al. Transcriptomic profiling reveals the molecular responses of *Rhodococcus aetherivorans* DMU1 to skatole stress. Ecotoxicol. Environ. Saf. 249, 114464 (2023).

22. Ma, Q., Zhang, X. & Qu, Y. Biodegradation and biotransformation of indole: Advances and perspectives. Front. Microbiol. 9, 2625 (2018).

23. Laird, T. S., Flores, N. & Leveau, J. H. J. Bacterial catabolism of indole-3-acetic acid. Appl. Microbiol. Biotechnol. 104, 9535–9550 (2020).

24. Farkas, C., Donoso, R. A., Melis-Arcos, F., Gárate-Castro, C. & Pérez-Pantoja, D. Complete genome sequence of *Rhodococcus ruber* R1, a novel strain showing a broad catabolic potential toward lignin-derived aromatics. Microbiol. Resour. Announc. 9, (2020).

25. Cappelletti, M. et al. Genome sequence of *Rhodococcus* sp. Strain BCP1, a biodegrader of alkanes and chlorinated compounds. Genome Announc. 1, (2013).

26. Haak, B., Fetzner, S. & Lingens, F. Cloning, nucleotide sequence, and expression of the plasmid-encoded genes for the two-component 2-halobenzoate 1,2-dioxygenase from *Pseudomonas cepacia* 2CBS. J. Bacteriol. 177, 667–675 (1995).

27. Eaton, R. W. & Chapman, P. J. Formation of indigo and related compounds from indolecarboxylic acids by aromatic acid-degrading bacteria: chromogenic reactions for cloning genes encoding dioxygenases that act on aromatic acids. J. Bacteriol. 177, 6983–6988 (1995).

28. Eaton, R. W. p-Cumate catabolic pathway in *Pseudomonas putida* Fl: cloning and characterization of DNA carrying the cmt operon. J. Bacteriol. 178, 1351–1362 (1996).

29. Kweon, O. et al. A new classification system for bacterial Rieske non-heme iron aromatic ring-hydroxylating oxygenases. BMC Biochem. 9, 11 (2008).

30. Barry, S. M. & Challis, G. L. Mechanism and catalytic diversity of Rieske non-heme iron-dependent oxygenases. ACS Catal. 3, 2362–2370 (2013).

31. Normand, P. et al. Genome characteristics of facultatively symbiotic *Frankia* sp. strains reflect host range and host plant biogeography. Genome Res. 17, 7–15 (2007).

32. Malone, K. M. et al. Updated reference genome sequence and annotation of *Mycobacterium bovis* AF2122/97. Genome Announc. 5, (2017).

33. Slesarev, A. I. et al. The complete genome of hyperthermophile *Methanopyrus kandleri* AV19 and monophyly of archaeal methanogens. Proc. Natl. Acad. Sci. U. S. A. 99, 4644–4649 (2002).

34. Ballut, L., Violot, S., Kumar, S., Aghajari, N. & Balaram, H. GMP synthetase: Allostery, structure, and function. Biomolecules 13, (2023).

35. Fujii, T., Takeo, M. & Maeda, Y. Plasmid-encoded genes specifying aniline oxidation from *Acinetobacter* sp. strain YAA. Microbiology 143 (Pt 1), 93–99 (1997).

36. Takeo, M. et al. Function of a glutamine synthetase-like protein in bacterial aniline oxidation via γ-glutamylanilide. J. Bacteriol. 195, 4406–4414 (2013).

37. Ji, J. et al. The substrate specificity of aniline dioxygenase is mainly determined by two of its components: glutamine synthetase-like enzyme and oxygenase. Appl. Microbiol. Biotechnol. 103, 6333–6344 (2019).

38. Li, S. et al. Complete biodegradation of fungicide carboxin and its metabolite aniline by *Delftia* sp. HFL-1. Sci. Total Environ. 912, 168957 (2024).

39. Oelschlägel, M., Zimmerling, J. & Tischler, D. A review: The styrene metabolizing cascade of side-chain oxygenation as biotechnological basis to gain various valuable compounds. Front. Microbiol. 9, 490 (2018).

40. Gassner, G. T. The styrene monooxygenase system. Methods Enzymol. 620, 423–453 (2019).

41. Sucharitakul, J., Chaiyen, P., Entsch, B. & Ballou, D. P. The reductase of p-hydroxyphenylacetate 3-hydroxylase from *Acinetobacter baumannii* requires p-hydroxyphenylacetate for effective catalysis. Biochemistry 44, 10434–10442 (2005).

42. Sucharitakul, J. et al. Kinetics of a two-component p-hydroxyphenylacetate hydroxylase explain how reduced flavin is transferred from the reductase to the oxygenase. Biochemistry 46, 8611–8623 (2007).

43. Marjanovic, A. et al. Catalytic and structural properties of ATP-dependent caprolactamase from *Pseudomonas jessenii*. Proteins 89, 1079–1098 (2021).

44. Kavanagh, K. L., Jörnvall, H., Persson, B. & Oppermann, U. Medium- and short-chain dehydrogenase/reductase gene and protein families : the SDR superfamily: functional and structural diversity within a family of metabolic and regulatory enzymes. Cell. Mol. Life Sci. 65, 3895–3906 (2008).

45. Sánchez-Cortés, S., Francioso, O., García-Ramos, J. V., Ciavatta, C. & Gessa, C. Catechol polymerization in the presence of silver surface. Colloids Surf. A Physicochem. Eng. Asp. 176, 177– 184 (2001).

46. Takeo, M., Fujii, T. & Maeda, Y. Sequence analysis of the genes encoding a multicomponent dioxygenase involved in oxidation of aniline and o-toluidine in *Acinetobacter* sp. strain YAA. J. Ferment. Bioeng. 85, 17–24 (1998).

47. Urata, M. et al. Genes involved in aniline degradation by *Delftia acidovorans* strain 7N and its distribution in the natural environment. Biosci. Biotechnol. Biochem. 68, 2457–2465 (2004).

48. Correll, C. C., Batie, C. J., Ballou, D. P. & Ludwig, M. L. Phthalate dioxygenase reductase: a modular structure for electron transfer from pyridine nucleotides to [2Fe-2S]. Science 258, 1604–1610 (1992).

49. Fukumori, F. & Saint, C. P. Nucleotide sequences and regulational analysis of genes involved in conversion of aniline to catechol in *Pseudomonas putida* UCC22(pTDN1). J. Bacteriol. 179, 399– 408 (1997).

50. Murakami, S., Hayashi, T., Maeda, T., Takenaka, S. & Aoki, K. Cloning and functional analysis of aniline dioxygenase gene cluster, from *Frateuria* species ANA-18, that metabolizes aniline via an ortho-cleavage pathway of catechol. Biosci. Biotechnol. Biochem. 67, 2351–2358 (2003).

51. Zhang, T., Zhang, J., Liu, S. & Liu, Z. A novel and complete gene cluster involved in the degradation of aniline by *Delftia* sp. AN3. J. Environ. Sci. (China) 20, 717–724 (2008).

52. Nagy, I. et al. Degradation of the thiocarbamate herbicide EPTC (S-ethyl dipropylcarbamothioate) and biosafening by *Rhodococcus* sp. strain NI86/21 involve an inducible cytochrome P-450 system and aldehyde dehydrogenase. J. Bacteriol. 177, 676–687 (1995).

53. Vieira, S., et al. *Capillimicrobium parvum* gen. nov., sp. nov., a novel representative of Capillimicrobiaceae fam. nov. within the order Solirubrobacterales, isolated from a grassland soil. Int. J. Syst. Evol. Microbiol. 72, (2022).

54. Walters, E. M. & Johnson, M. K. Ferredoxin:Thioredoxin reductase: Disulfide reduction catalyzed via novel site-specific [4Fe-4S] cluster chemistry. Photosynth. Res. 79, 249–264 (2004).

55. Jacquot, J.-P., Eklund, H., Rouhier, N. & Schürmann, P. Structural and evolutionary aspects of thioredoxin reductases in photosynthetic organisms. Trends Plant Sci. 14, 336–343 (2009).

56. Prakash, D. et al. Toward a mechanistic and physiological understanding of a ferredoxin:disulfide reductase from the domains Archaea and Bacteria. J. Biol. Chem. 293, 9198–9209 (2018).

57. Zhang, C., Freddolino, P. L. & Zhang, Y. COFACTOR: improved protein function prediction by combining structure, sequence and protein-protein interaction information. Nucleic Acids Res. 45, W291–W299 (2017).

58. Liu, Q., Yang, L.-L. & Xin, Y.-H. Diversity of the genus *Cryobacterium* and proposal of 19 novel species isolated from glaciers. Front. Microbiol. 14, 1115168 (2023).

59. Siddiqi, M. Z., Lee, S.-Y., Yeon, J. M. & Im, W.-T. *Arthrobacter hankyongi* sp. nov., Isolated From Wet Land. Curr. Microbiol. 80, 92 (2023).

60. Kummer, C., Schumann, P. & Stackebrandt, E. *Gordonia alkanivorans* sp. nov., isolated from tar-contaminated soil. Int. J. Syst. Bacteriol. 49 Pt 4, 1513–1522 (1999).

61. Chaturvedi, N. K. & Katoch, S. S. Remedial technologies for aniline and aniline derivatives elimination from wastewater. J. Health Pollut. 10, 200302 (2020).

62. Radomski, J. L. The primary aromatic amines: their biological properties and structure-activity relationships. Annu. Rev. Pharmacol. Toxicol. 19, 129–157 (1979).

63. Benigni, R. & Passerini, L. Carcinogenicity of the aromatic amines: from structure-activity relationships to mechanisms of action and risk assessment. Mutat. Res. 511, 191–206 (2002).

64. Ferraz, E. R. A. The impact of aromatic amines on the environment risks and damages. Front. Biosci. (Elite Ed.) E4, 914–923 (2012).

65. Donoso, R. A., González-Toro, F. & Pérez-Pantoja, D. Widespread distribution of hmf genes in Proteobacteria reveals key enzymes for 5-hydroxymethylfurfural conversion. Comput. Struct. Biotechnol. J. 19, 2160–2169 (2021).

66. Schmittgen, T. D. & Livak, K. J. Analyzing real-time PCR data by the comparative C(T) method. Nat. Protoc. 3, 1101–1108 (2008).

67. Gibson, D. G. et al. Enzymatic assembly of DNA molecules up to several hundred kilobases. Nat. Methods 6, 343–345 (2009).

68. Donoso, R. A., Corbinaud, R., Gárate-Castro, C., Galaz, S. & Pérez-Pantoja, D. Identification of a phylogenetically divergent vanillate O-demethylase from *Rhodococcus ruber* R1 supporting growth on meta-methoxylated aromatic acids. Microorganisms 11, 78 (2022).

69. Seemann, T. Prokka: rapid prokaryotic genome annotation. Bioinformatics 30, 2068–2069 (2014).

70. Page, A. J. et al. Roary: rapid large-scale prokaryote pan genome analysis. Bioinformatics 31, 3691– 3693 (2015).

71. NCBI Resource Coordinators et al. Database resources of the National Center for Biotechnology Information. Nucleic Acids Res. 46, D8–D13 (2018).

72. Katoh, K., Rozewicki, J. & Yamada, K. D. MAFFT online service: multiple sequence alignment, interactive sequence choice and visualization. Brief. Bioinform. 20, 1160–1166 (2019).

73. Nguyen, L.-T., Schmidt, H. A., von Haeseler, A. & Minh, B. Q. IQ-TREE: a fast and effective stochastic algorithm for estimating maximum-likelihood phylogenies. Mol. Biol. Evol. 32, 268–274 (2015).

74. Letunic, I. & Bork, P. Interactive Tree of Life (iTOL) v6: recent updates to the phylogenetic tree display and annotation tool. Nucleic Acids Res. 52, W78–W82 (2024).

75. Wang, J. et al. The conserved domain database in 2023. Nucleic Acids Res. 51, D384–D388 (2023).

76. Jumper, J. et al. Highly accurate protein structure prediction with AlphaFold. Nature 596, 583–589 (2021).

