## Supplementary information, Galaz et al 2024 for "Aniline Dioxygenase in *Rhodococcus ruber* R1: Insights into Skatole Degradation"

Supplementary Tables S1-S2

Supplementary Figures S1-S8

### Supplementary Table S1

**Table S1.** Bacterial strains and plasmids used in this study.

| Strain or plasmid | Relevant phenotype and/or genotype | Reference or source |
|---|---|---|
| ***Rhodococcus* strains** | | |
| *R. ruber* R1 | $Skt^+$, $Aniline^-$, $Bz^+$, $Fructose^+$ | 1 |
| *R. ruber* Chol-4 | $Skt^-$, $Aniline^-$, $Bz^+$, $Fructose^+$ | 2 |
| *R. ruber* DSM 43338[T] | $Skt^-$, $Aniline^-$, $Bz^+$, $Fructose^+$ | DSMZ[a] |
| *R. aetherivorans* BCP1 | $Skt^+$, $Aniline^-$, $Bz^+$, $Fructose^+$ | 3 |
| **Other strains** | | |
| *E. coli* Mach1 | ΔrecA1398 endA1 tonAΦ80ΔlacM15 ΔlacX74 hsdR ($rK^-mK^+$) | Invitrogen, Carlsbad, USA |
| *G. alkanivorans* NBRC 16433 | $SKt^-$, $Aniline^+$, $Fructose^+$ | NBRC[b] |
| *C. pinatubonensis* JMP134 | $Skt^-$, $Aniline^-$, $Bz^+$, $Fructose^+$ | 4 |
| *C. pinatubonensis* JMP222 | $Skt^-$, $Aniline^-$, $Bz^+$, $Fructose^+$ | 5 |
| **Plasmids** | | |
| pBS1 | Broad host range vector, *araC-$P_{BAD}$*, $Gm^R$ | 6 |
| pBS1-*sktF* | pBS1 derivative expressing *sktF*genes, $Gm^R$ | This study |
| pBS1-*sktA-F* | pBS1 derivative expressing *sktABZCDEF* genes, $Gm^R$ | This study |

a: DSMZ, Deutsche Sammlung von Mikroorganismen und Zellkulturen GmbH (Braunschweig, Germany).

b: NBRC, Biological Resource Center, NITE.

T: Type strain.

Abbreviations: Skt, skatole; Bz, Benzoate

### Supplementary Table S2

**Table 2.** Primer pairs used in this study.

| Purpose | Forward primer | Sequence (5' – 3') | Reverse primer | Sequence (5' – 3') |
|---|---|---|---|---|
| **Real-Time PCR** | | | | |
| | 16S-Fw-RT | CTGGACATAAGGGGCATGAT | 16S-Rv-RT | CTGTGTTGCCAGCACGTAAT |
| | sktA-Fw-RT | GAGGTGTTCGAGGAGCTGAT | sktA-Rv-RT | AGGTTCGGGAAGATGTTGAG |
| | sktG-Fw-RT | ACCGGCAAGTACGGATTGT | sktG-Rv-RT | CCGAAGAGGTTGAAGCACAC |
| | sktJ-Fw-RT | GGAGAACTCCTCGGGATCTG | sktJ-Rv-RT | GCACGTTGATCTCGTACCAC |
| | sktL-Fw-RT | AACTCGCTGACCTCCCTGT | sktL-Rv-RT | CACGTGGACTGCGAAATACT |
| | RS07680-Fw-RT | GATCGCGACGATCAATCTG | RS07680-Rv-RT | ATCATCGTGGTGAGGTTGGT |
| | RS07690-Fw-RT | AATCCGGTATCGGAAGACG | RS07690-Rv-RT | GTGTCCTTCCCGACCACTC |
| | RS07770-Fw-RT | TCCTGGTCTACGAGAACACCT | RS07770-Rv-RT | AGATCCAAGAGCGTCCAGAG |
| | RS07790-Fw-RT | CACCTGCACTACACCACGAC | RS07790-Rv-RT | GTAGACGTGCACGAGGTTGA |
| | catA1-Fw-RT | ACAACCGCAAGCACTTCTC | catA1-Rv-RT | ATGGCCCTTGAAGATCAGC |
| | catA2-Fw-RT | ACCTGCCGCAGTGGAATC | catA2-Rv-RT | CGTCGGGATCTGGTACGG |
| | benA-Fw-RT | GACAAGACCGAGGTCACCAT | benA-Rv-RT | GGAACTCCTCGAGATCGTCA |
| | rps7-Fw-RT | CCAGCTGGTCAACAAGATCC | rps7-Rv-RT | ATCGGTGCCGGTCTTCTC |
| **Expression constructs** | | | | |
| | sktA-Fw1 | TTGGGCTAGCGAATTCCTGCGAA AGGGCAAAAGCCATGA | sktC-Rv1 | GAACTCCGGTGCCTTGTC |
| | sktC-Fw2 | ACGAAATCCGGGACAAGG | sktF-Rv2 | GTAATACGACTCACTATAGGGA GCGGTGGTCAGAAGAGTT |
| | sktF-Fw3 | TTGGGCTAGCGAATTCCTGCCGT CTTCTTCAGTTCGGTGC | sktF-Rv3 | GTAATACGACTCACTATAGGAG CGGTGGTCAGAAGAGTTC |
| | pBS1a | GCAGGAATTCGCTAGCCCAA | pBS1b | CCTATAGTGAGTCGTATTAC |

### Supplementary Figure S1

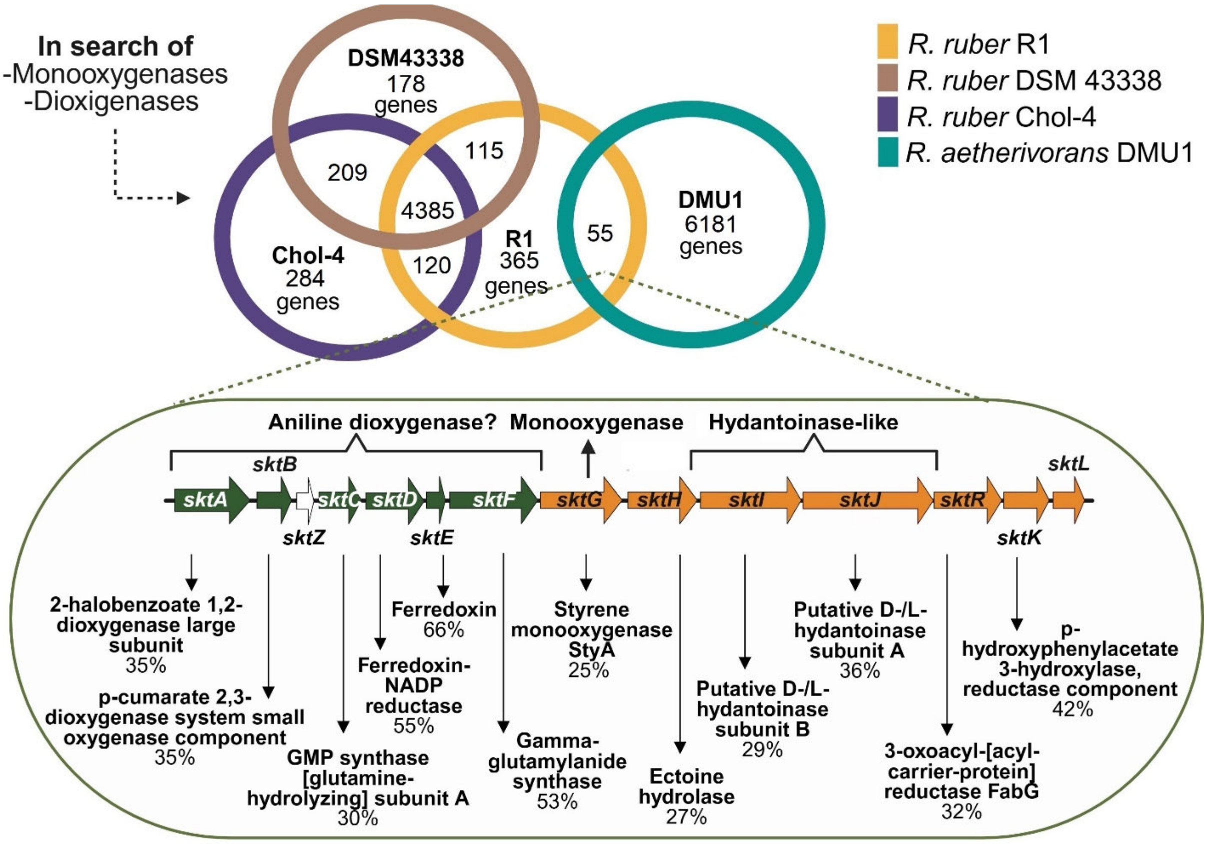

**Figure S1. Identification of the potential genetic cluster involved in skatole degradation by *Rhodococcus ruber* R1 strain.** Comparative genomics of the R1 strain, to other closely related *R. ruber* strains that do not degrade skatole (Chol-4 and DSM43338), as well as the skatole-degrading strain *R. aetherivorans* DMU1, confirmed an unusual set of 14 genes previously identified in DMU1 (Li *et al.*, 2023), which we refer to as *skt* genes. The figure depicts genes encoding putative aniline dioxygenase (green arrows), monoxygenase activity, and hydantoinase-like enzymes. Additionally, percentages of amino acid identities with proteins of demonstrated function were obtained for each translated protein present in the skatole cluster through a search in the UniProtKB/Swiss-Prot database. A hypothetical protein, named SktZ (indicated by a white arrow), was identified on the putative aniline dioxygenase encoding genes.

**Supplementary Figure S2**

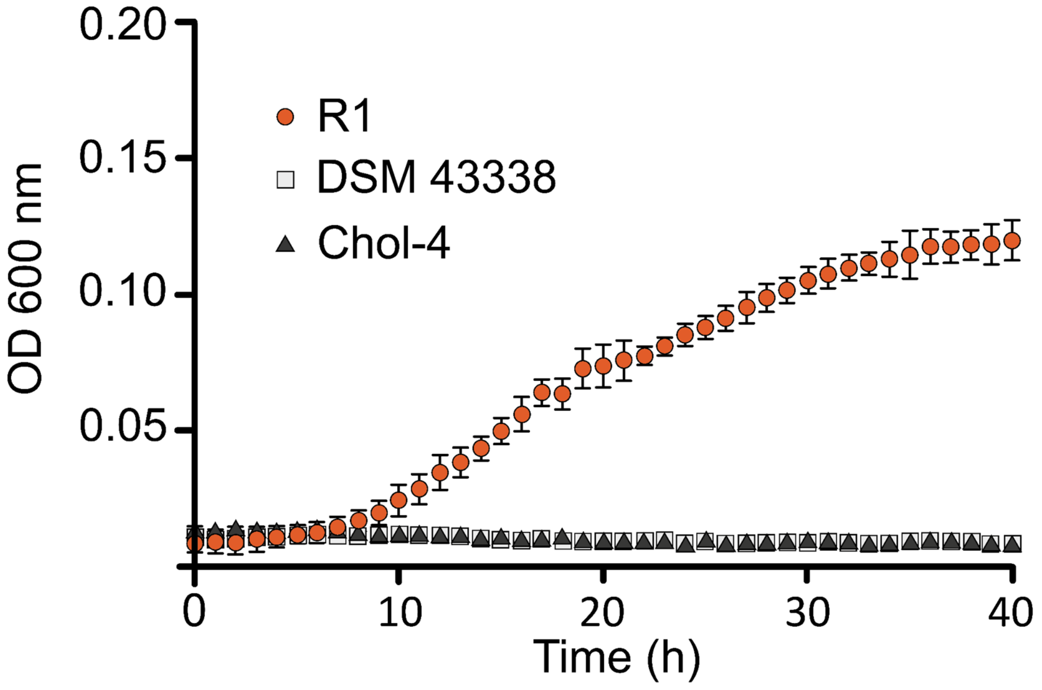

**Figure S2. Additional *Rhodococcus ruber* strains cannot grow on skatole.** Strains Chol-4, DSM43338, and R1 were grown in mineral salt medium with 1 mM skatole as the sole carbon and energy source. Experiments were performed in three biological replicates. Error bars: ±SD.

**Supplementary Figure S3**

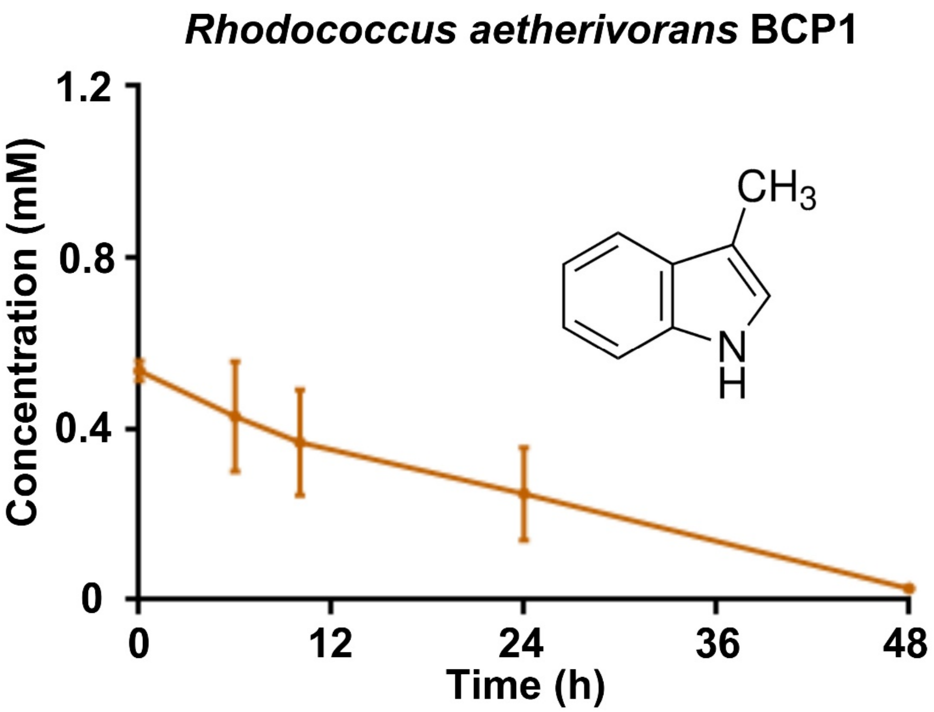

**Figure S3. *Rhodococcus aetherivorans* BCP1 degrades skatole.** Cells of *Rhodococcus aetherivorans* BCP1 were grown on 20 mM fructose. Cells were washed and then exposed to 0.5 mM skatole. Three biological replicates were performed for substrate consumption measurements. Error bars: ±SD.

### Supplementary Figure S4

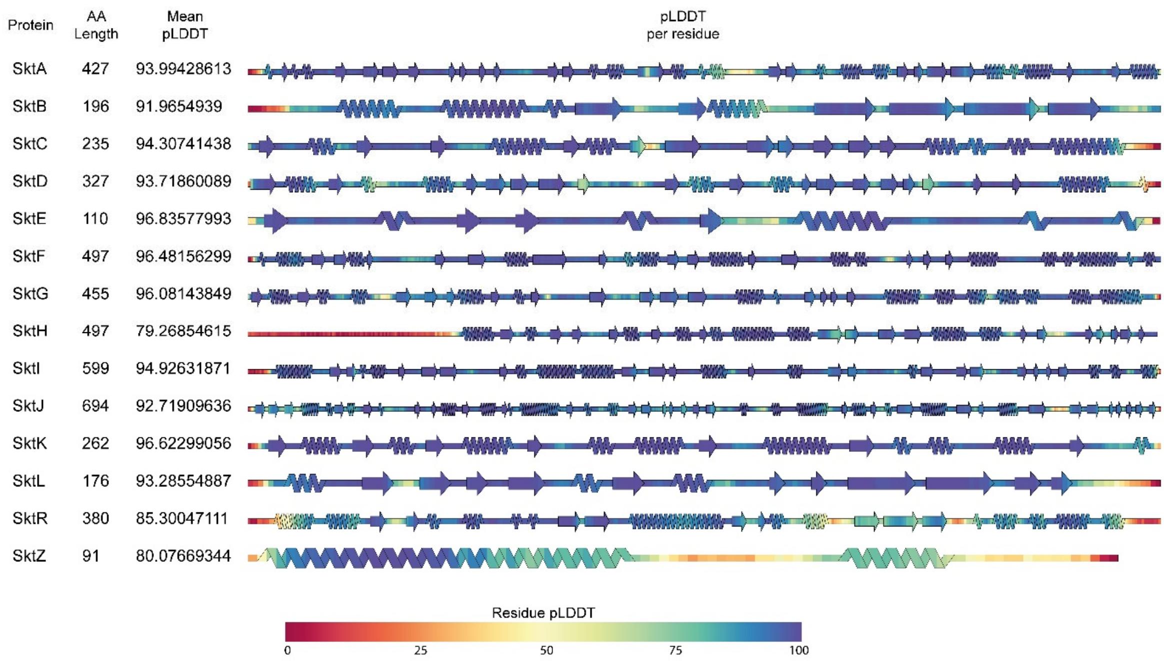

**Figure S4. Assessment of Protein Structure Reliability Based on pLDDT Scores.** Protein structure prediction using AlphaFold (Jumper *et al.*, 2021) as implemented in AlphaFold2 available on Google Colab, configured with 3 recycles and a maximum of 200 iterations, excluding Amber relaxation. The optimal model for each protein was selected based on the pLDDT score, which quantifies the confidence of AlphaFold2 in its predictions. The selected PDB files were subsequently processed with SSDraw, where the secondary structure was color-coded according to individual pLDDT values, with scores approaching 100 indicating higher reliability. To enhance clarity, the figures were aligned and standardized in width.

### Supplementary Figure S5

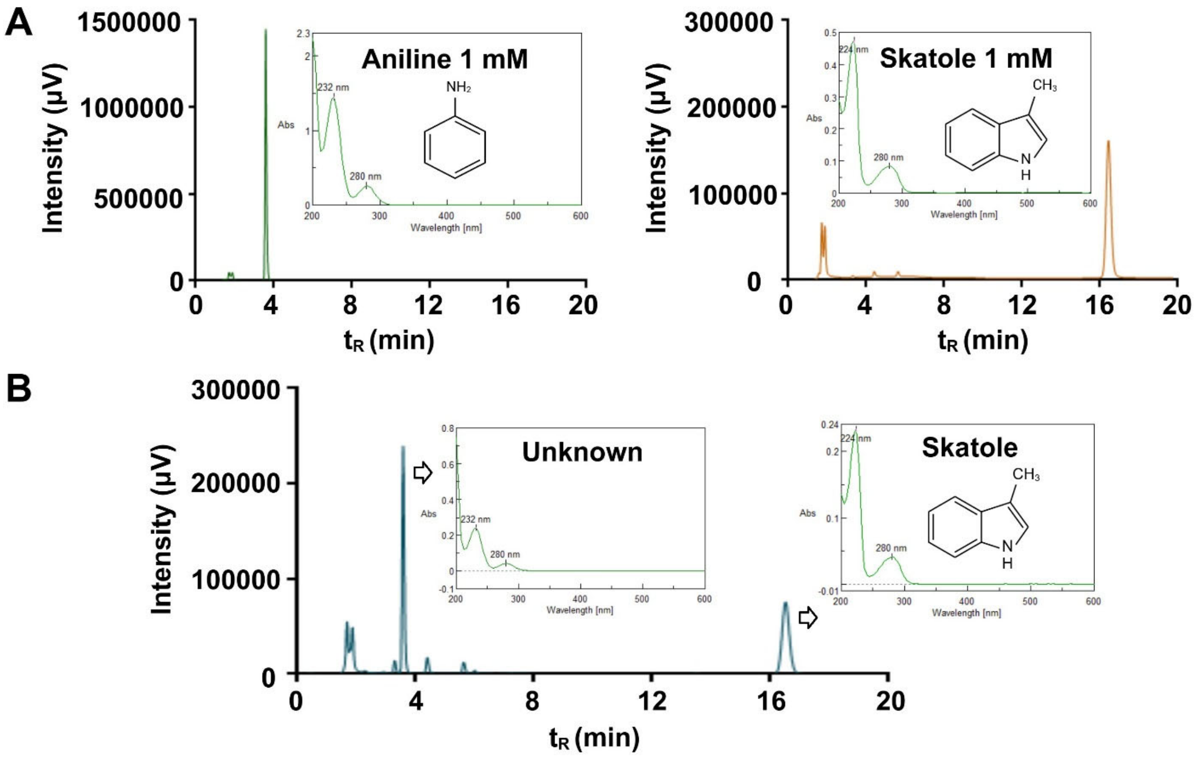

**Figure S5**. **Identification of aniline as a possible intermediate in the degradation of skatole.** (A) Chromatograms and absorption spectrum of 1 mM aniline and skatole standards. (B) The chromatogram and absorption spectrum of the supernatant were analyzed at 27 hours during the growth of strain R1 in skatole (see Figure 2), revealing the appearance of an unknown compound at 3.5 minutes, concurrent with skatole (retention time 16.5 minutes).

### Supplementary Figure S6

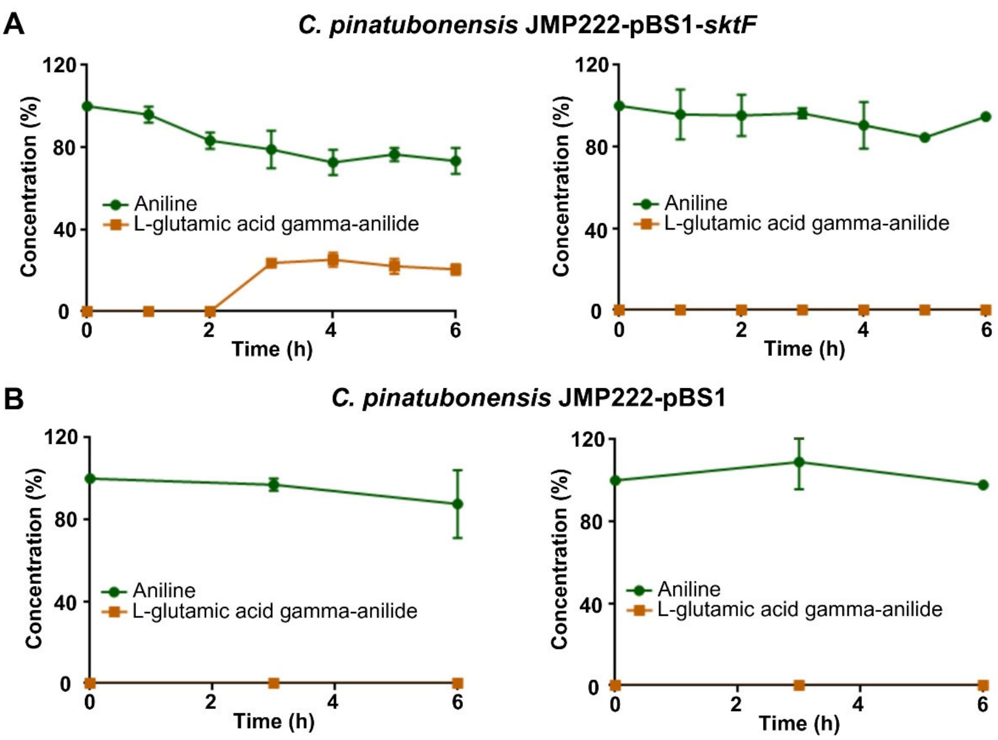

**Figure S6. Protein SktF functions as a γ-GA synthetase.** (A) Cells of *C. pinatubonensis* JMP222 expressing sktF gene or (B) carrying the empty vector pBS1 (control) were grown on 20 mM fructose supplemented with 2,5 mM arabinose as an inducer for 3 h. The cells were then washed and subsequently exposed to 1 mM aniline and 1 mM glutamate (left graph) or only 1 mM aniline (right graph) for 6h. Three biological replicates were performed to determine the percentage degradation. Error bars: ±SD.

### Supplementary Figure S7

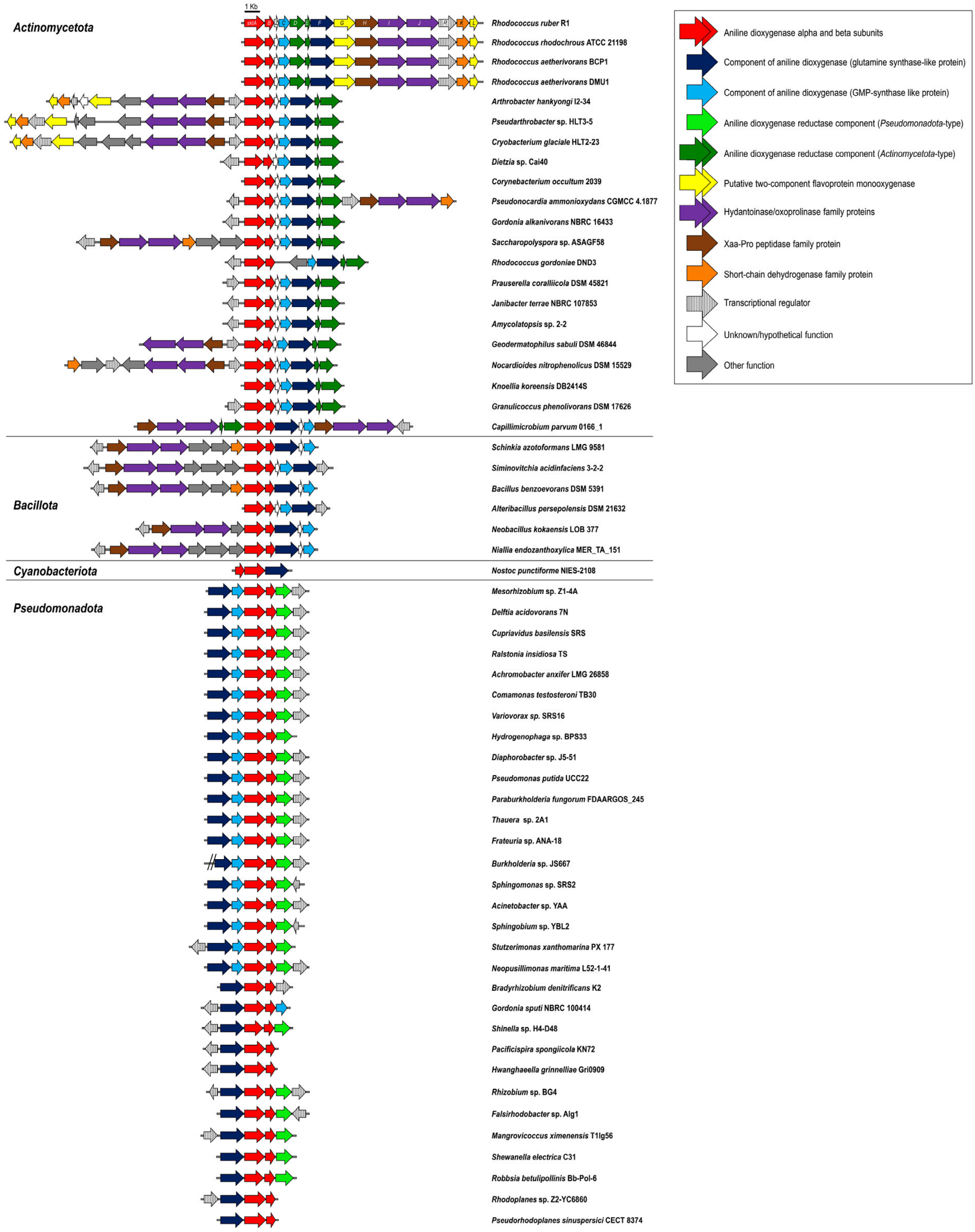

Actinomycetota
1 Kb
sktA
B
Z
C
D
E
F
G
H
I
J
R
K
L
Rhodococcus ruber R1
Rhodococcus rhodochrous ATCC 21198
Rhodococcus aetherivorans BCP1
Rhodococcus aetherivorans DMU1
Arthrobacter hankyongi I2-34
Pseudarthrobacter sp. HLT3-5
Cryobacterium glaciale HLT2-23
Dietzia sp. Cai40
Corynebacterium occultum 2039
Pseudonocardia ammonioxydans CGMCC 4.1877
Gordonia alkanivorans NBRC 16433
Saccharopolyspora sp. ASAGF58
Rhodococcus gordoniae DND3
Prauserella coralliicola DSM 45821
Janibacter terrae NBRC 107853
Amycolatopsis sp. 2-2
Geodermatophilus sabuli DSM 46844
Nocardioides nitrophenolicus DSM 15529
Knoellia koreensis DB2414S
Granulicoccus phenolivorans DSM 17626
Capillimicrobium parvum 0166_1
Bacillota
Schinkia azotoformans LMG 9581
Siminovitchia acidinfaciens 3-2-2
Bacillus benzoevorans DSM 5391
Alteribacillus persepolensis DSM 21632
Neobacillus kokaensis LOB 377
Niallia endozanthoxylica MER_TA_151
Cyanobacteriota
Nostoc punctiforme NIES-2108
Pseudomonadota
Mesorhizobium sp. Z1-4A
Delftia acidovorans 7N
Cupriavidus basilensis SRS
Ralstonia insidiosa TS
Achromobacter anxifer LMG 26858
Comamonas testosteroni TB30
Variovorax sp. SRS16
Hydrogenophaga sp. BPS33
Diaphorobacter sp. J5-51
Pseudomonas putida UCC22
Paraburkholderia fungorum FDAARGOS_245
Thauera sp. 2A1
Frateuria sp. ANA-18
Burkholderia sp. JS667
Sphingomonas sp. SRS2
Acinetobacter sp. YAA
Sphingobium sp. YBL2
Stutzerimonas xanthomarina PX 177
Neopusillimonas maritima L52-1-41
Bradyrhizobium denitrificans K2
Gordonia sputi NBRC 100414
Shinella sp. H4-D48
Pacificispira spongiicola KN72
Hwanghaeella grinnelliae Gri0909
Rhizobium sp. BG4
Falsirhodobacter sp. Alg1
Mangrovicoccus ximenensis T1lg56
Shewanella electrica C31
Robbsia betulipollinis Bb-Pol-6
Rhodoplanes sp. Z2-YC6860
Pseudorhodoplanes sinuspersici CECT 8374
Aniline dioxygenase alpha and beta subunits
Component of aniline dioxygenase (glutamine synthase-like protein)
Component of aniline dioxygenase (GMP-synthase like protein)
Aniline dioxygenase reductase component (Pseudomonadota-type)
Aniline dioxygenase reductase component (Actinomycetota-type)
Putative two-component flavoprotein monooxygenase
Hydantoinase/oxoprolinase family proteins
Xaa-Pro peptidase family protein
Short-chain dehydrogenase family protein
Transcriptional regulator
Unknown/hypothetical function
Other function

**Figure S7. Clusters of skatole and aniline degradation genes in representative bacterial species.** The conserved *sktA* gene of strain R1 was used to align the *skt* gene clusters from selected species of Pseudomonadota, Actinomycetota, Bacillota, and Cyanobacteryota lineages. Genes that are putatively encoding enzymes and regulators of the biodegradative pathway for skatole/aniline are colored or shaded according to their functions, as depicted in the figure.

**Supplementary Figure S8**

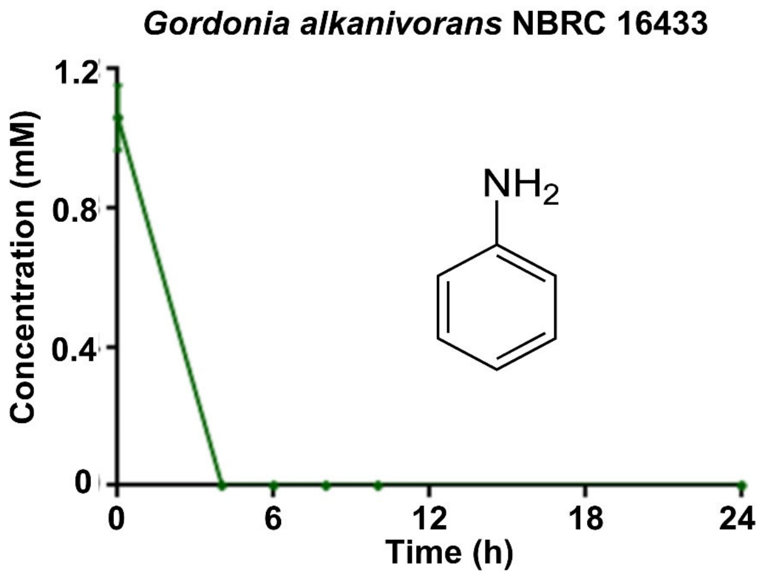

**Figure S8. *Gordonia alkanivorans* NBRC 16433 degrades aniline.** Resting cells of *Gordonia alkanivorans* NBRC 16433 were grown on 20 mM fructose induced with 0.5 mM aniline, were washed and then exposed to 1 mM aniline. Three biological replicates were performed for substrate consumption measurements. Error bars: ±SD.

### REFERENCES

1. Farkas, C., Donoso, R. A., Melis-Arcos, F., Gárate-Castro, C. & Pérez-Pantoja, D. Complete genome sequence of *Rhodococcus ruber* R1, a novel strain showing a broad catabolic potential toward lignin-derived aromatics. *Microbiol. Resour. Announc.* **9**, (2020).
2. Fernández de Las Heras, L., García Fernández, E., María Navarro Llorens, J., Perera, J. & Drzyzga, O. Morphological, physiological, and molecular characterization of a newly isolated steroid-degrading actinomycete, identified as *Rhodococcus ruber* strain Chol-4. Curr Microbiol. **59**, 548–553 (2009).
3. Cappelletti, M. *et al.* Genome sequence of *Rhodococcus* sp. Strain BCP1, a biodegrader of alkanes and chlorinated compounds. *Genome Announc.* **1**, (2013).
4. Pérez-Pantoja, D., De la Iglesia, R., Pieper, D.H., and González, B. Metabolic reconstruction of aromatic compounds degradation from the genome of the amazing pollutant-degrading bacterium *Cupriavidus necator* JMP134. FEMS Microbiol Rev. **32**: 736–794 (2008).
5. Padilla, L., Matus, V., Zenteno, P., and González, B. Degradation of 2,4,6-trichlorophenol via chlorohydroxyquinol in *Ralstonia eutropha* JMP134 and JMP222. J Basic Microbiol. **40**: 243–249 (2000).
6. Donoso, R.A., Corbinaud, R., Gárate-Castro, C., Galaz, S., and Pérez-Pantoja, D. Identification of a phylogenetically divergent vanillate O-demethylase from *Rhodococcus ruber* R1 supporting growth on meta-methoxylated aromatic acids. Microorganisms. **11**: 78 (2022).
